# Reduced Gene Dosage of Histone H4 Prevents CENP-A Mislocalization in *Saccharomyces cerevisiae*

**DOI:** 10.1101/2020.05.29.124032

**Authors:** Jessica R. Eisenstatt, Kentaro Ohkuni, Wei-Chun Au, Olivia Preston, Evelyn Suva, Michael Costanzo, Charles Boone, Munira A. Basrai

## Abstract

Mislocalization of the centromeric histone H3 variant (Cse4 in budding yeast, CID in flies, CENP-A in humans) to non-centromeric regions contributes to chromosomal instability (CIN) in yeast, fly, and human cells. Overexpression and mislocalization of CENP-A has been observed in cancers, however, the mechanisms that facilitate the mislocalization of overexpressed CENP-A have not been fully explored. Defects in ubiquitin-mediated proteolysis of overexpressed Cse4 (*GALCSE4*) leads to its mislocalization and synthetic dosage lethality (SDL) in mutants for E3 ubiquitin ligases (Psh1, Slx5, SCF^Met30^, SCF^Cdc4^), Doa1, Hir2, and Cdc7. In contrast, defects in sumoylation of *GALcse4K215/216/A/R* prevent its mislocalization and do not cause SDL in a *psh1*Δ strain. Here, we used a genome-wide screen to identify factors that facilitate the mislocalization of overexpressed Cse4 by characterizing suppressors of the *psh1*Δ *GALCSE4* SDL. Deletions of histone H4 alleles (*HHF1* or *HHF*2), which were among the most prominent suppressors, also suppress *slx5Δ, cdc4-1, doa1Δ, hir2Δ*, and *cdc7-4 GALCSE4* SDL. Reduced dosage of *H4* contributes to defects in sumoylation and reduced mislocalization of overexpressed Cse4. We determined that the *hhf1-20, cse4-102*, and *cse4-111* mutants, which are defective in the Cse4-H4 interaction, also exhibit reduced sumoylation of Cse4 and do not display *psh1*Δ *GALCSE4* SDL. In summary, we have identified genes that contribute to the mislocalization of overexpressed Cse4 and defined a role for the gene dosage of *H4* in facilitating Cse4 sumoylation and mislocalization to non-centromeric regions, contributing to SDL when Cse4 is overexpressed in mutant strains.

## INTRODUCTION

Centromeres are specialized chromosome loci that are essential for faithful chromosome segregation during mitosis and meiosis. The kinetochore (centromeric DNA and associated proteins) provides an attachment site for microtubules to promote proper segregation of sister chromatids during cell division (Allshire and Karpen 2008; Verdaasdonk and Bloom 2011; Burrack and Berman 2012; Choy *et al.* 2012; Maddox *et al.* 2012; McKinley and Cheeseman 2016). Despite the wide divergence of centromeric DNA sequence, establishment of centromeric chromatin is regulated by epigenetic mechanisms where incorporation of the essential and evolutionarily conserved centromeric histone H3 variant CENP-A (Cse4 in *Saccharomyces cerevisiae*, Cnp1 in *Schizosaccharomyces pombe*, CID in *Drosophila melanogaster*, and CENP-A in mammals) serves to nucleate kinetochore assembly (Kitagawa and Hieter 2001; Biggins 2013; McKinley and Cheeseman 2016).

The evolutionarily conserved CENP-A-specific histone chaperones (Scm3 in *S. cerevisiae* and *S. pombe*, CAL1 in *D. melanogaster*, Holliday Junction Recognition Protein HJURP in humans) mediate the centromeric localization of CENP-A (Camahort *et al.* 2007; Mizuguchi *et al.* 2007; Stoler *et al.* 2007; Foltz *et al.* 2009; Pidoux *et al.* 2009; Williams *et al.* 2009; Shuaib *et al.* 2010; Chen *et al.* 2014). In budding yeast, other chaperones such as Chromatin Assembly Factor 1 (CAF-1), an evolutionarily conserved replication-coupled histone H3/H4 chaperone, can facilitate the deposition of overexpressed Cse4 when Scm3 is depleted (Hewawasam et al. 2018). The CAF-1 orthologues Mis16 in *S. pombe* and RbAp46/48 in humans and *D. melanogaster* also contribute to centromeric localization of CENP-A (Fujita *et al.* 2007; Pidoux *et al.* 2009; Williams *et al.* 2009; Boltengagen *et al.* 2016).

Restricting the localization of CENP-A to centromeres is essential for faithful chromosome segregation. However, overexpression of CENP-A leads to its mislocalization to non-centromeric chromatin and contributes to chromosomal instability (CIN) in yeast, flies, and humans (Collins *et al.* 2004; Heun *et al.* 2006; Moreno-Moreno *et al.* 2006; Au *et al.* 2008; Mishra *et al.* 2011; Lacoste *et al.* 2014; Athwal *et al.* 2015; Shrestha *et al.* 2017). Overexpression and mislocalization of CENP-A is observed in many cancers and is proposed to promote tumorigenesis (Tomonaga *et al.* 2003; Amato *et al.* 2009; Li *et al.* 2011; McGovern *et al.* 2012; Sun *et al.* 2016). Thus, defining the molecular mechanisms that promote and prevent mislocalization of CENP-A is an area of active investigation.

In budding yeast, post-translational modifications (PTMs) of Cse4, such as ubiquitination, sumoylation, and isomerization, are important for regulating steady-state levels of Cse4 and preventing its mislocalization to non-centromeric regions, thereby maintaining chromosome stability (Collins *et al.* 2004; Hewawasam *et al.* 2010; Ranjitkar *et al.* 2010; Ohkuni *et al.* 2014; Ohkuni *et al.* 2016; Cheng *et al.* 2017; Au *et al.* 2020). Ubiquitin-mediated proteolysis of Cse4 by E3 ubiquitin ligases such as Psh1 (Hewawasam *et al.* 2010; Ranjitkar *et al.* 2010), SUMO-targeted ubiquitin ligase (STUbL) Slx5 (Ohkuni *et al.* 2016), SCF^Met30/Cdc4^ (Au *et al.* 2020), SCF^Rcy1^ (Cheng *et al.* 2016), and Ubr1 (Cheng *et al.* 2017) and the proline isomerase Fpr3 (Ohkuni *et al.* 2014) regulate the cellular levels of Cse4. Psh1-mediated proteolysis of Cse4 has been well characterized and has been shown to be regulated by the FACT (Facilitates Chromatin Transcription/Transactions) complex (Deyter and Biggins 2014), CK2 (Casein Kinase 2) (Hewawasam *et al.* 2014), HIR (HIstone Regulation) histone chaperone complex (Ciftci-Yilmaz *et al.* 2018), and DDK (Dbf4-Dependent Kinase) complex (Eisenstatt *et al.* 2020). In general, mutation or deletion of factors that prevent Cse4 mislocalization show synthetic dosage lethality (SDL) when Cse4 is overexpressed from a galactose-inducible promoter (*GALCSE4*).

In contrast to the many studies that have characterized pathways that prevent mislocalization of CENP-A to non-centromeric regions, mechanisms that facilitate the mislocalization of overexpressed CENP-A have not been fully explored. Studies from our laboratory and those of others show that the transcription-coupled histone H3/H4 chaperone DAXX/ATRX promotes mislocalization of CENP-A to non-centromeric regions in human cells (Lacoste *et al.* 2014; Shrestha *et al.* 2017). In budding yeast, CAF-1 contributes to the mislocalization of overexpressed Cse4 to non-centromeric regions (Hewawasam *et al.* 2018). We have recently shown that sumoylation of Cse4K215/216 in the C-terminus of Cse4 facilitates its interaction with CAF-1 and this promotes the deposition of Cse4 to non-centromeric regions (Ohkuni *et al.* 2020). Notably, *psh1*Δ *cac2*Δ *GALCSE4* strains and *psh1*Δ *GALcse4K215/216R/A* strains do not exhibit SDL due to reduced mislocalization of Cse4 (Hewawasam *et al.* 2018; Ohkuni *et al.* 2020).

Defining the mechanisms that facilitate the mislocalization of overexpressed Cse4 to non-centromeric regions is essential for understanding which pathways contributes to mislocalization of CENP-A in cancers with a poor prognosis. We performed a genome-wide screen using a synthetic genetic array (SGA) to identify genes that promote Cse4 mislocalization. We took advantage of the SDL of a *psh1Δ GALCSE4* strain (Hewawasam *et al.* 2010; Ranjitkar *et al.* 2010; Au *et al.* 2013) to identify suppressors of the SDL phenotype. An SGA analysis was performed by combining mutants of essential genes and deletion of non-essential genes with *psh1*Δ *GALCSE4*. The screen identified mutations or deletions of genes encoding regulators of chromatin remodeling, RNA transcription/processing, nucleosome occupancy, ubiquitination, and histone H4. The budding yeast genome possesses two gene pairs which encode almost identical H3 and H4 proteins (*HHT1/HHF1* and *HHT2/HHF2*) and two gene pairs which encode identical H2A and H2B proteins (*HTA1/HTB1* and *HTA2/HTB2*). Deletion of the two alleles that encode histone H4 (*HHF1* or *HHF2*) were among the most prominent suppressors of the *psh1Δ GALCSE4* SDL. A role for the dosage of *H4* in preventing mislocalization of Cse4 has not been previously examined.

In this study, we focused on defining the molecular mechanisms that prevent the mislocalization of overexpressed Cse4 and suppress the *psh1Δ GALCSE4* SDL when the gene dosage of *H4* is reduced. We showed that deletion of *HHF1* or *HHF2* also suppresses the *GALCSE4* SDL in *slx5Δ, doa1Δ, hir2Δ, cdc4-1*, and *cdc7-4* strains. Deletion of *HHF1* or *HHF2* results in reduced Cse4 sumoylation and this correlates with reduced mislocalization to non-centromeric regions and rapid degradation of Cse4 in a *psh1Δ* strain. Moreover, *cse4-102, cse4-111*, and *hhf1-20*, which have mutations in their histone fold domains and are defective for the formation of the Cse4-H4 dimer (Smith *et al.* 1996; Glowczewski *et al.* 2000), show reduced Cse4 sumoylation and do not cause SDL in *psh1Δ GALCSE4* strains. In summary, our genome-wide suppressor screen allowed us to identify genes that contribute to Cse4 mislocalization and to define a role for reduced gene dosage of *H4* in preventing the mislocalization of Cse4 to non-centromeric regions and suppression of the *psh1Δ GALCSE4* SDL.

## MATERIALS AND METHODS

### Strains and plasmids

Yeast strains used in this study are described in Table S2 and plasmids in Table S3. Yeast strains were grown in rich media (1% yeast extract, 2% bacto-peptone, 2% glucose) or synthetic medium with glucose or raffinose and galactose (2% final concentration each) and supplements to allow for selection of the indicated plasmids. Double mutant strains were generated by mating wild type or *psh1Δ* strains with empty vector or a plasmid containing *GAL1-6His-3HA-CSE4* to mutant strains on rich medium at room temperature for six hours followed by selection of diploid cells on medium selective for the plasmid and appropriate resistance markers. Diploids were sporulated for 5 days at 23°C and plated on selective medium without uracil, histidine, or arginine and with canavanine, clonNAT, and G418 to select for *MAT*a double mutants. The synthetic genetic array (SGA) was performed as previously described (Costanzo *et al.* 2016).

### Growth assays

Growth assays were performed as previously described (Eisenstatt *et al.* 2020). Wild type and mutant strains were grown on medium selective for the plasmid, suspended in water to a concentration with an optical density of 1 measured at a wavelength of 600 nm (OD_600_, approximately 1.0 × 10^7^ cells per ml), and plated in five-fold serial dilutions starting with 1 OD_600_ on synthetic growth medium containing glucose or galactose and raffinose (2% final concentration each) selecting for the plasmid. Strains were grown at the indicated temperatures for 3-5 days.

### Protein stability assays

Protein stability assays were performed as previously described (Au *et al.* 2008). Briefly, logarithmically growing wild type and mutant cells were grown for three to four hours in media selective for the plasmid containing galactose/raffinose (2% final concentration each) at 30°C followed by addition of cycloheximide (CHX, 10 µg/ml) and glucose (2% final concentration). Protein extracts were prepared from cells collected 0, 30, 60, and 90 minutes after CHX addition with the TCA method as described previously (Kastenmayer *et al.* 2006). Equal amount of protein as determined by the Bio-Rad DC™ Protein Assay were analyzed by Western blot. Proteins were separated by SDS-PAGE on 4-12% Bis-TRIS SDS-polyacrylamide gels (Novex, NP0322BOX) and analysis was done against primary antibodies α-HA (1:1000, Roche, 12CA5) or α-Tub2 (1:4500, custom made for Basrai Laboratory) in TBST containing 5% (w/v) dried skim milk. HRP-conjugated sheep α-mouse IgG (Amersham Biosciences, NA931V) and HRP-conjugated donkey α-rabbit IgG (Amersham Biosciences, NA934V) were used as secondary antibodies. Stability of the Cse4 protein relative to the Tub2 loading control was measured as the percent remaining as determined with the Image Lab Software (BioRad).

### Ubiquitination pull-down assay

Levels of ubiquitinated Cse4 were determined with ubiquitin pull-down assays as described previously (Au *et al.* 2013) with modifications. Cells were grown to logarithmic phase, induced in galactose-containing medium for 3 hours at 30°C and pelleted. The cell pellet was resuspended in lysis buffer (20 mM Na_2_HPO_4_, 20 mM NAH_2_PO_4_, 50 mM NaF, 5 mM tetra-sodium pyrophosphate, 10 mM beta-glycerolphosphate, 2 mM EDTA, 1 mM DTT, 1% NP-40, 5 mM N-Ethylmaleimide, 1 mM PMSF, and protease inhibitor cocktail (Sigma, catalogue # P8215)) and equal volume of glass beads (lysing matrix C, MP Biomedicals). Cell lysates were generated by homogenizing cells with a FastPrep-24 5G homogenizer (MP Biomedicals) and a fraction of the lysate was aliquoted for input. An equal concentration of lysates from wild type and mutant strains were incubated with tandem ubiquitin binding entities (Agarose-TUBE1, Life Sensors, Inc., catalogue # UM401) overnight at 4°C. Proteins bound to the beads were washed three times with TBS-T at room temperature and eluted in 2 x Laemmli buffer at 100°C for 10 minutes. The eluted protein was resolved on a 4-12% Bis-Tris gel (Novex, NP0322BOX) and ubiquitinated Cse4 was detected by Western blot using anti-HA antibody (Roche Inc., 12CA5). Levels of ubiquitinated Cse4 relative to the non-modified Cse4 in the input were quantified using software provided by the Syngene imaging system. The percentage of ubiquitinated Cse4 levels is set to 100% in the wild type strain.

### *In vivo* sumoylation assay

Cell lysates were prepared from 50 ml culture of strains grown to logarithmic phase in raffinose/galactose (2% final concentration each) medium at 30°C for 4 hours to induce expression of Cse4 from the galactose-inducible promoter. Cells were pelleted, rinsed with sterile water, and suspended in 0.5 ml of guanidine buffer (0.1 M Tris-HCl at pH 8.0, 6.0 M guanidine chloride, 0.5 M NaCl). Cells were homogenized with Matrix C (MP Biomedicals) using a bead beater (MP Biomedicals, FastPrep-24 5G). Cell lysates were clarified by centrifugation at 6,000 rpm for 7 min and protein concentration was determined using a DC protein assay kit (Bio-Rad). Samples containing equal amounts of protein were brought to a total volume of 1 ml with appropriate buffer.

*In vivo* sumoylation was assayed in crude yeast extracts using nickel-nitrilotriacetic acid (Ni-NTA) agarose beads to pull down His-HA-tagged Cse4 as described previously (Ohkuni *et al.* 2015) with modifications. Cell lysates were incubated with 100 μl of Ni-NTA superflow beads (Qiagen, 30430) overnight at 4 °C. After being washed with guanidine buffer one time and with breaking buffer (0.1 M Tris-HCl at pH 8.0, 20 % glycerol, 1 mM PMSF) five times, beads were incubated with 2x Laemmli buffer including imidazole at 100°C for 5 min. The protein samples were analyzed by SDS-PAGE and western blotting. Primary antibodies were anti-HA (12CA5) mouse (Roche, 11583816001) and anti-Smt3 (y-84) rabbit (Santa Cruz Biotechnology, sc-28649). Secondary antibodies were ECL Mouse IgG, HRP-Linked Whole Ab (GE Healthcare Life Sciences, NA931V) or ECL Rabbit IgG, HRP-linked Whole Ab (GE Healthcare Life Sciences, NA934V). Protein levels were quantified using Image Lab software (version 6.0.0) from Bio-Rad Laboratories, Inc (Hercules, CA).

### ChIP-qPCR

Chromatin immunoprecipitations were performed with two biological replicates per strain as previously described (Cole *et al.* 2014; Chereji *et al.* 2017; Eisenstatt *et al.* 2020) with modifications. Logarithmic phase cultures were grown in raffinose/galactose (2% final concentration each) media for 4 hours and were treated with formaldehyde (1% final concentration) for 20 minutes at 30°C followed by the addition of 2.5 M glycine for 10 minutes at 30°C. Cell pellets were washed twice with 1 X PBS and resuspended in 2 mL FA Lysis Buffer (1 mM EDTA pH8.0, 50 mM HEPES-KOH pH7.5, 140 mM NaCl, 0.1% sodium deoxycholate, 1% Triton X-100) with 1 x protease inhibitors (Sigma) and 1 mM PMSF (final concentration). The cell suspension was split into four screw top tubes with glass beads (0.4-0.65 mm diameter) and lysed in a FastPrep-24 5G (MP Biosciences) for 40 seconds three times, allowed to rest on ice for 5 minutes, and lysed two final times for 40 seconds each. The cell lysate was collected, and the chromatin pellet was washed in FA Lysis Buffer twice. Each pellet was resuspended in 600 µl of FA Lysis Buffer and combined into one 5 ml tube. The chromatin suspension was sonicated with a Branson digital sonifer 24 times at 20% amplitude with a repeated 15 seconds on/off cycle. After 3 minutes of centrifugation (13000 rpm, 4°C), the supernatant was transferred to another tube. Input sample was removed (5%) and the average size of the DNA was analyzed. The remaining lysate was incubated with anti-HA-agarose beads (Sigma, A2095) overnight at 4°C. The beads were washed in 1 ml FA, FA-HS (500 mM NaCl), RIPA, and TE buffers for five minutes on a rotor two times each. The beads were suspended in ChIP Elution Buffer (25 mM Tris-HCl pH7.6, 100 mMNaCl, 0.5% SDS) and incubated at 65°C overnight. The beads were treated with proteinase K (0.5 mg/ml) and incubated at 55°C for four hours followed by Phenol/Chloroform extraction and ethanol precipitation. The DNA pellet was resuspended in a total of 50 µl sterile water. Samples were analyzed by quantitative PCR (qPCR) performed with the 7500 Fast Real Time PCR System with Fast SYBR Green Master Mix (Applied Biosystems). qPCR conditions used: 95°C for 20 sec; 40 cycles of 95°C for 3 sec, 60° for 30 sec. Primers used are listed in Table S4.

### Data availability

Strains and plasmids are available upon request. Supporting figures S1-S7 are available as JPG files. Supporting Table S1 is an Excel file that describes mutations that suppress the *psh1Δ GALCSE4* SDL, the gene systematic name, the gene name, the functional category, growth and colony scores, and validation information if applicable. File S1 contains Tables S2, S3, and S4 which describe the yeast strains, plasmids, and primers used in this study, respectively. Supporting information is available at FigShare.

## RESULTS

### A genome-wide screen identified suppressors of the SDL in a *psh1Δ GALCSE4* strain

Identifying pathways that facilitate the deposition of overexpressed Cse4 to non-centromeric regions will provide insight into the mechanisms that promote chromosomal instability (CIN) in CENP-A overexpressing cancers. Deletion of *PSH1*, which regulates ubiquitin-mediated proteolysis of overexpressed Cse4, results in synthetic dosage lethality (SDL) when Cse4 is overexpressed (*GALCSE4*) (Hewawasam *et al.* 2010; Ranjitkar *et al.* 2010). We reasoned that strains with deletions or mutations of factors that promote Cse4 mislocalization would rescue the SDL of a *psh1Δ GALCSE4* strain. Therefore, we generated a *psh1Δ* query strain overexpressing *CSE4* from a galactose-inducible plasmid and mated it to arrays of 3,827 non-essential gene deletion strains and of 786 conditional mutant alleles, encoding 560 essential genes, and 186 non-essential genes for internal controls (Costanzo *et al.* 2016). Growth of the haploid meiotic progeny plated in quadruplicate was visually scored on glucose-and galactose-containing media grown at 30°C for non-essential and 26°C for essential gene mutant strains (Figure 1A). Highlighted in the figure are all four replicates of deletion of histone H4 (*hhf1Δ*) and Hap3 (*hap3Δ*) showing better growth on galactose media compared to the control strains along the perimeter and other deletion strains on the plate (Figure 1B, bottom and top square, respectively). Strains that suppress the *psh1Δ GALCSE4* SDL on galactose-containing media were given a growth score of one (low suppression) to four (high suppression) (Table S1). The number of replicates within the quadruplicate that displayed the same growth were given a colony score of one (one out of four replicates) to four (all four replicates). We identified ninety-four deletion and mutant alleles encoding ninety-two genes that suppressed the *psh1Δ GALCSE4* SDL and the majority (81%) of quadruplicates had all four colonies displaying the same level of suppression, indicated by a colony score of four (Table S1).

**Figure 1.**
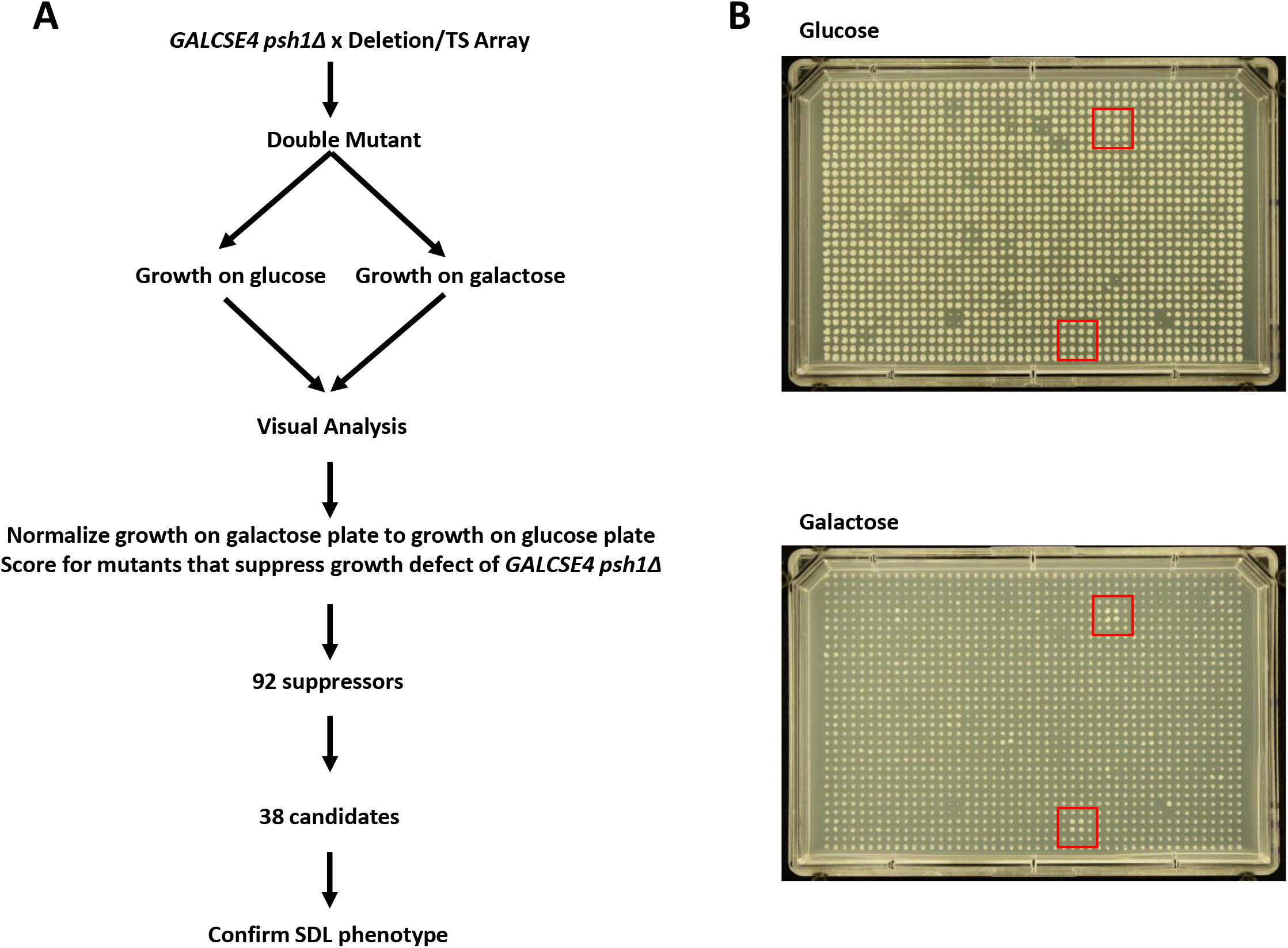
A genome-wide screen identified suppressors of the *psh1Δ GALCSE4* SDL. **A. Schematic for the genome-wide screen.** A *psh1Δ* strain (YMB8995) transformed with *GAL1-6His-3HA-CSE4* (pMB1458) was mated to an array of non-essential gene deletions and an array of conditional alleles of essential genes. Growth of the haploid meiotic progeny plated in quadruplicate was visually scored on glucose-and galactose-containing media grown at 30°C for non-essential and 26°C for essential gene mutant strains. Ninety-two genes were identified as growing better on galactose-containing media than the *psh1Δ GALCSE4* strain. Thirty-eight candidate genes were selected for confirmation of suppression of lethality. **B. Representative plates from the genome-wide screen.** Shown is Plate 01 of the non-essential gene deletion array. The mutant strains were spotted in quadruplicate on selective media plates containing glucose (top) or galactose (bottom). Red boxes (top box is *hap3Δ*; bottom box is *hhf1Δ*) highlight mutant strains that displayed improved growth on galactose-containing plates compared to the *psh1Δ GALCSE4* control strain (perimeter of plate) and did not show a growth defect or improved growth on the glucose plates.

Of the ninety-four alleles, we selected thirty-eight candidate mutants (fourteen non-essential deletion strains and twenty-four conditional mutants) to confirm the suppression of the *psh1Δ GALCSE4* SDL (Table 1). These candidates displayed a growth score of three or four where most of the replicates displayed high suppression and represent pathways involved in RNA processing and cleavage, DNA repair, chromatin remodeling, histone modifications, and DNA replication (Table 1). Secondary validation of the SDL suppressors was done by independently generating double mutant strains of *psh1Δ GALCSE4* with candidate mutants. Growth assays were performed on media selective for the *GALCSE4* plasmid and containing either glucose or raffinose and galactose. We used a *hir2Δ psh1Δ* strain as a negative control because *hir2Δ psh1Δ GALCSE4* strains display SDL (Ciftci-Yilmaz *et al.* 2018). Of the thirty-eight strains tested, twenty-nine showed almost complete suppression, five strains showed a partial suppression, and four did not suppress the SDL on galactose media (Tables 1 and S1 and Figures S1A and S1B). We further tested a subset of the thirty-eight genes to confirm overexpression of *CSE4* and found that strains with mutations in genes involved in RNA processing and transcription do not show galactose-induced expression of *CSE4* (Table S1 and Figure S1C), indicating that these are false positive hits. Through secondary validation, we confirmed that 89% of the candidate mutants tested suppressed the *psh1Δ GALCSE4* SDL.

**TABLE 1.**
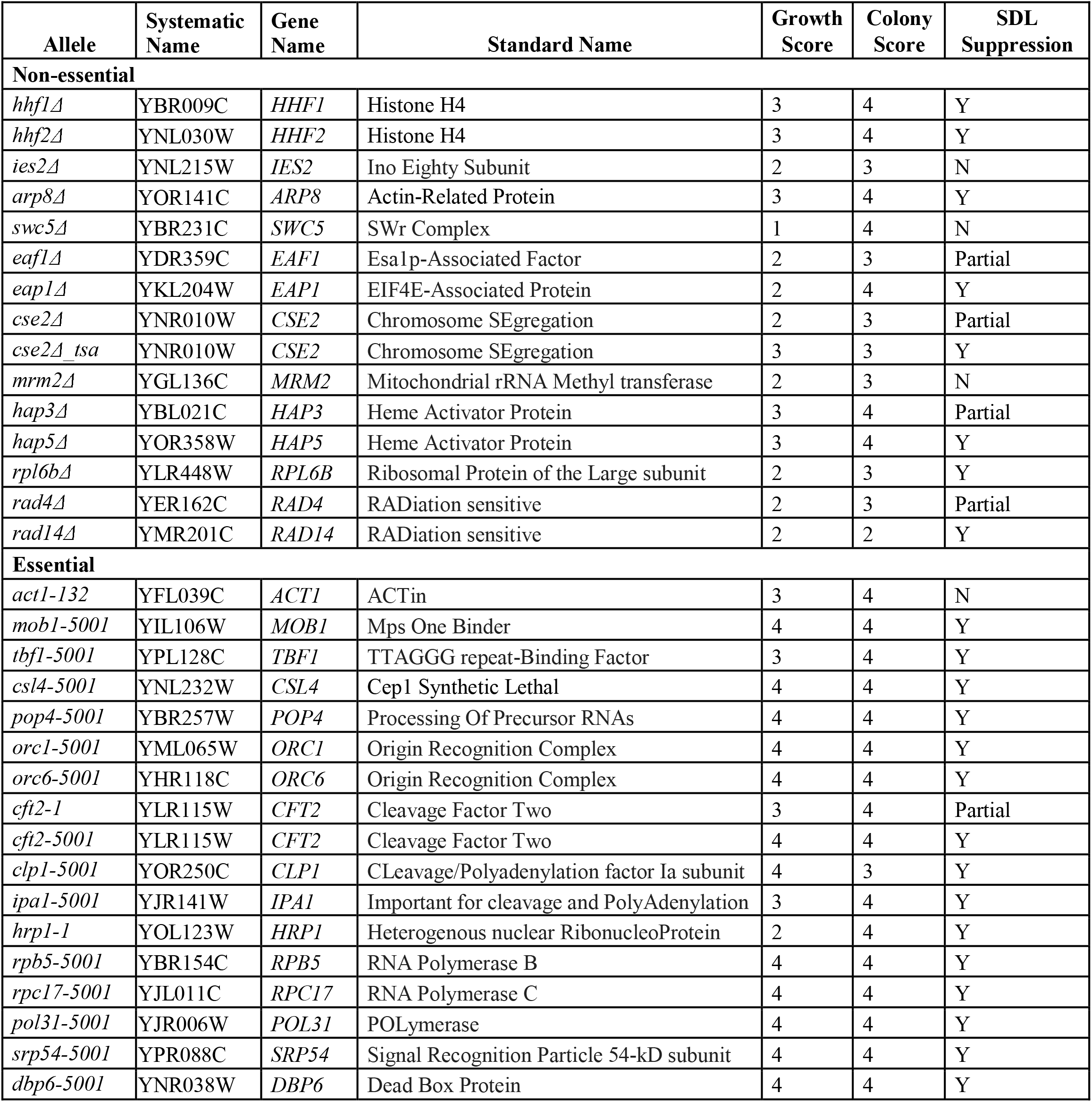
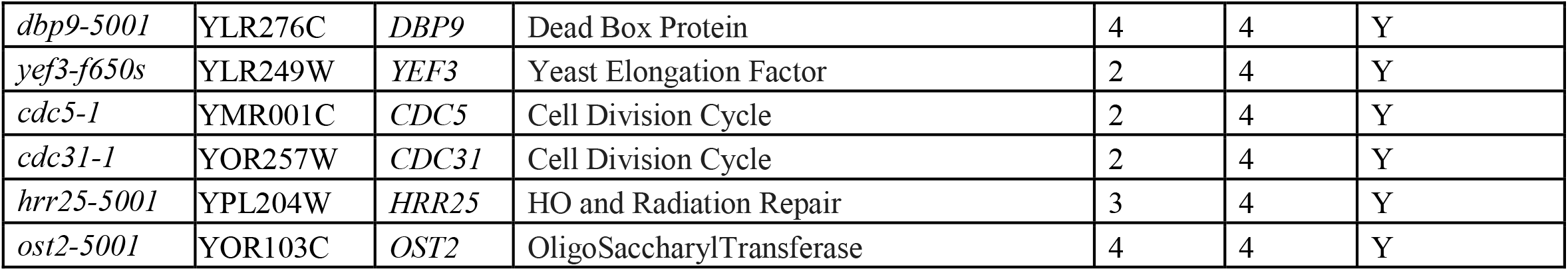
Candidate double mutant strains with the indicated mutant allele combined with *psh1Δ GALCSE4* were generated and used for secondary validation using growth assays. Indicated is the allele analyzed, systematic name, gene name, standard name, visual scoring from the primary screen for growth score (from one to four) and colony score (from one to four), and suppression of SDL (Y: SDL was suppressed; N: SDL was not suppressed; Partial: SDL was partially suppressed).

We initiated our studies with the INO80 chromatin remodeling complex as our screen identified deletion and mutant alleles corresponding to three components of the INO80 complex, Ies2, Arp8, and Act1 (Poch and Winsor 1997; Shen *et al.* 2000; Shen *et al.* 2003; Tosi *et al.* 2013). Secondary validation assays showed that *ies2Δ* and *act1-132* do not suppress the SDL of the *psh1Δ GALCSE4* strain (Figures S1A and S1B). In contrast, *arp8Δ* did suppress the *psh1Δ GALCSE4* SDL (Figures S1A and S2A) however, the *arp8Δ* strain displayed polyploidy when analyzed by Fluorescent Activated Cell Sorting (FACS) (Figure S2B) and we consequently did not pursue further studies with the INO80 complex.

### Deletion of histone H4 alleles suppresses the SDL of a *psh1Δ GALCSE4* strain

Two nonallelic loci, *HHT1/HHF1* and *HHT2/HHF2*, encode identical H3 and H4 proteins in budding yeast. The screen identified the deletion of either one of the histone H4 alleles, *HHT1/hhf1*Δ (*hhf1Δ*) or *HHT2/hhf2*Δ (*hhf2Δ*), as among the most prominent suppressors of the *psh1Δ GALCSE4* SDL. A role for the dosage of histone H4-encoding genes in mislocalization of Cse4 has not yet been reported. We confirmed that the *hhf1Δ* and *hhf2Δ* strains do not exhibit defects in ploidy or cell cycle by FACS analysis (Figure S3). Growth assays confirmed that *psh1Δ hhf1Δ GALCSE4* and *psh1Δ hhf2Δ GALCSE4* strains plated on galactose media do not exhibit SDL (Figure 2A). We determined that the phenotype was linked to deletion of the H4 alleles because transformation of a plasmid with the respective wild type histone H4 gene into the *psh1Δ hhf1Δ* or *psh1Δ hhf2Δ* strains restored the SDL observed in the *psh1Δ GALCSE4* strain (Figure 2B).

**Figure 2.**
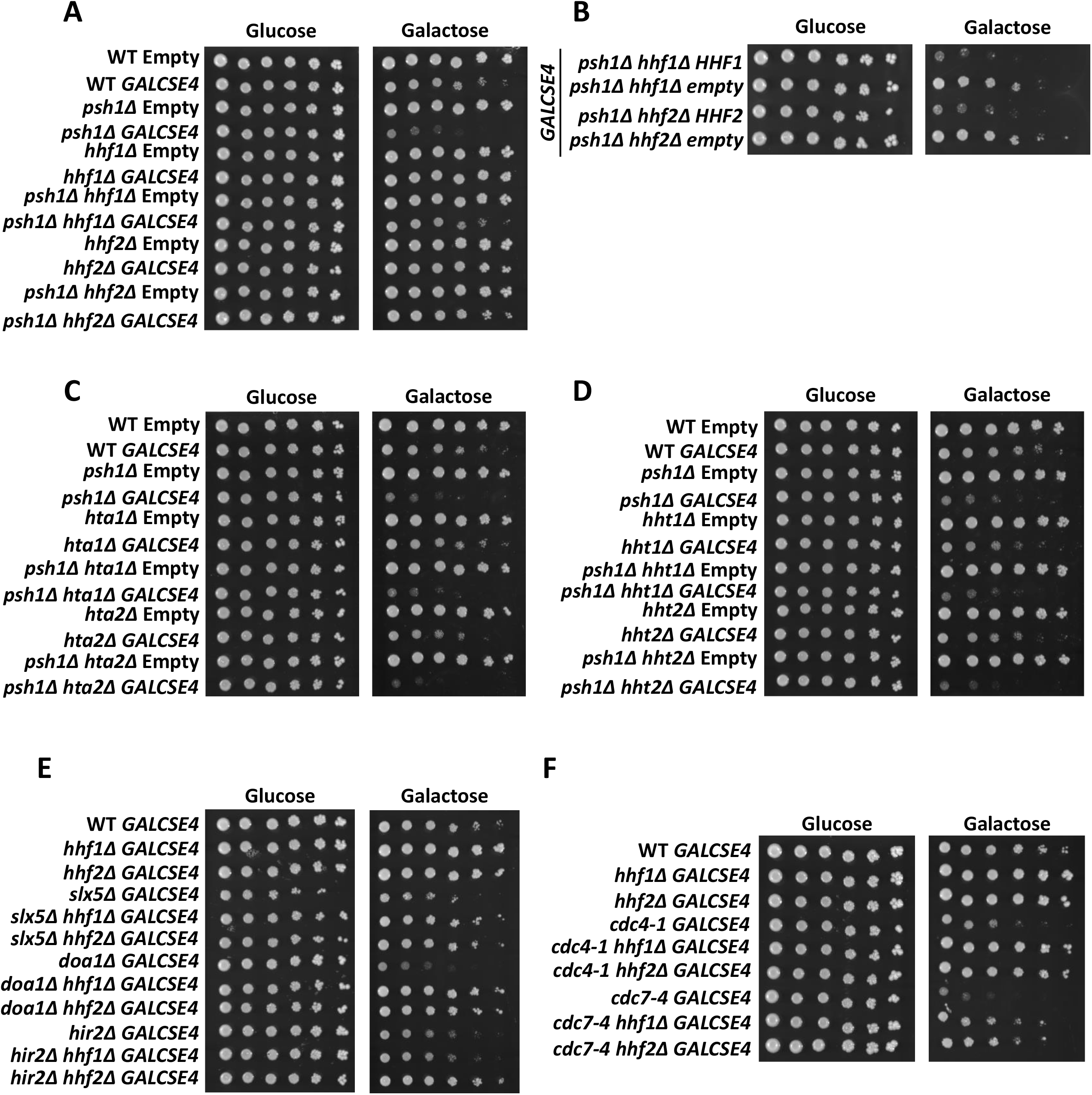
Deletion of *H4* genes suppresses *GALCSE4* SDL. Three independent isolates for each strain were assayed and shown is a representative for each. **A. The *psh1Δ GALCSE4* SDL is suppressed by deletion of *HHF1* or *HHF2*.** Growth assays of wild type, *psh1Δ, hhf1Δ, hhf2Δ, psh1Δ hhf1Δ*, and *psh1Δ hhf2Δ* strains with empty vector (pMB433; YMB9802, YMB10478, YMB10825, YMB11166, YMB10821, and YMB10823, respectively) or *GAL1-6His-3HA-CSE4* (pMB1458; YMB9803, YMB10479, YMB10937, YMB10938, YMB10822, and YMB10824, respectively). Cells were spotted in five-fold serial dilutions on glucose (2% final concentration) or raffinose/galactose (2% final concentration each) media selective for the plasmid and grown at 30°C for three to five days. **B. The *psh1Δ GALCSE4* SDL suppression is linked to the *hhf1Δ* and *hhf2Δ* alleles.** Growth assays of *psh1Δ hhf1Δ* (YMB10822) and *psh1Δ hhf2Δ* (YMB10824) strains with *GAL1-6His-3HA-CSE4* (pMB1458) transformed with empty vector (pRS425) or a plasmid containing wild type *HHF1* (pMB1928) or *HHF2* (pMB1929). Strains were assayed as described above in (A). **C.** and **D. Deletion of genes encoding histones H2A (C) or H3 (D) does not suppress the SDL of a *psh1Δ GALCSE4* strain.** Growth assays of wild type, *psh1Δ*, and (C) *hta1Δ, hta2Δ, psh1Δ hta1Δ, psh1Δ hta2Δ*, or (D) *hht1Δ, hht2Δ, psh1Δ hht1Δ*, and *psh1Δ hht1Δ* strains with empty vector (pMB433; YMB9802, YMB10478, YMB11258, YMB11266, YMB11260, YMB11268, YMB11274, YMB11282, YMB11276, and YMB11284, respectively) or *GAL1-6His-3HA-CSE4* (pMB1458: YMB9803, YMB10479, YMB11262, YMB11270, YMB11264, YMB11272, YMB11278, YMB11286, YMB11280, and YMB11288, respectively). Strains were assayed as described above in (A). **E. Reduced gene dosage of *H4* suppresses the SDL of *slx5Δ, doa1Δ*, and *hir2Δ GALCSE4* strains.** Growth assays of wild type (YMB9804), *hhf1Δ* (YMB10937), *hhf2Δ* (YMB10938), *slx5Δ* (YMB10963), *slx5Δ hhf1Δ* (YMB11046), *slx5Δ hhf2Δ* (YMB11047), *doa1Δ* (YMB11032), *doa1Δ hhf1Δ* (YMB11050), *doa1Δ hhf2Δ* (YMB11053), *hir2Δ* (YMB8332), *hir2Δ hhf1Δ* (YMB11105), *hir2Δ hhf2Δ* (YMB11107) strains expressing *GAL1-6HIS-3HA-CSE4* (pMB1458). Strains were assayed as described above in (A) and grown at 30°C for three to five days. **F. Deletion of *HHF1* or *HHF2* suppresses the SDL of *cdc4-1* and *cdc7-4 GALCSE4* strains.** Growth assays of wild type (YMB9804), *hhf1Δ* (YMB10937), *hhf2Δ* (YMB10938), *cdc4-1* (YMB9756), *cdc4-1 hhf1Δ* (YMB11051), *cdc4-1 hhf2Δ* (YMB11054), *cdc7-4* (YMB9760), *cdc7-4 hhf1Δ* (YMB11052), and *cdc7-4 hhf2Δ* (YMB11055) with *GAL1-6His-3HA-CSE4* (pMB1458). Strains were assayed as described above in (A) and grown at 23°C for three to five days.

We next investigated if deletion of a single allele for either histone H3 or H2A genes could suppress the SDL of a *psh1Δ GALCSE4* strain. Note that the two nonallelic loci, *HTA1/HTB1* and *HTA2/HTB2*, encode almost identical H2A and H2B proteins. Deletion of *HTA1* (*hta1*Δ*/HTB1*), *HTA2* (*hta2*Δ*/HTB2*), *HHT1* (*hht1*Δ*/HHF1*), or *HHT2* (*hht2*Δ*/HHF2*) did not suppress the SDL of a *psh1Δ GALCSE4* strain on galactose media (Figures 2C and 2D and Table 2). Based on these results we conclude that the suppression of *psh1Δ GALCSE4* SDL is specific to the reduced gene dosage of *H4*.

**TABLE 2.** Summary of the SDL growth phenotypes of mutants that exhibit SDL with *GALCSE4* and combined with *hhf1Δ* or *hhf2Δ*. Shown is the protein function, relevant strain genotype, and growth with *GALCSE4*. Wild type growth is indicated as ++; SDL as --- and extent of suppression (++ or +++).

| Protein Function | Relevant Strain Genotype | Growth with *GALCSE4* |
| --- | --- | --- |
|  | WT | ++ |
| Histone H4 | *hhf1Δ* | +++ |
|  | *hhf2Δ* | +++ |
| Histone H2A | *hta1Δ* | ++ |
|  | *hta2Δ* | ++ |
| Histone H3 | *hht1Δ* | ++ |
|  | *hht2Δ* | ++ |
| E3 Ubiquitin Ligase | *psh1Δ* | --- |
|  | *psh1Δ hhf1Δ* | +++ |
|  | *psh1Δ hhf2Δ* | +++ |
|  | *psh1Δ hhf1Δ + HHF1* | --- |
|  | *psh1Δ hhf2Δ + HHF2* | --- |
|  | *psh1Δ hta1Δ* | --- |
|  | *psh1Δ hta2Δ* | --- |
|  | *psh1Δ hht1Δ* | --- |
|  | *psh1Δ hht2Δ* | --- |
| SUMO-Targeted Ubiquitin Ligase | *slx5Δ* | -- |
|  | *slx5Δ hhf1Δ* | +++ |
|  | *slx5Δ hhf2Δ* | +++ |
| Ubiquitin Binding | *doa1Δ* | --- |
|  | *doa1Δ hhf1Δ* | +++ |
|  | *doa1Δ hhf2Δ* | +++ |
| HIR Nucleosome Binding Complex | *hir2Δ* | -- |
|  | *hir2Δ hhf1Δ* | - |
|  | *hir2Δ hhf2Δ* | ++ |
| F-box of the SCF Complex | *cdc4-1* | -- |
|  | *cdc4-1 hhf1Δ* | +++ |
|  | *cdc4-1 hhf2Δ* | +++ |
| Dbf4-Dependent Kinase | *cdc7-4* | --- |
|  | *cdc7-4 hhf1Δ* | ++ |
|  | *cdc7-4 hhf2Δ* | ++ |

### Reduced gene dosage of *H4* suppresses the SDL of *slx5Δ, doa1Δ, hir2Δ, cdc4-1*, and *cdc7-4 GALCSE4* strains

To determine if the SDL suppression by reduced *H4* gene dosage is limited to the *psh1Δ GALCSE4* strain, we deleted *HHF1* or *HHF2* in deletion or mutant strains encoding Slx5, Doa1, Hir2, Cdc4, and Cdc7 as deletion or mutation of these factors show SDL with *GALCSE4* and mislocalization of transiently overexpressed Cse4 (Au *et al.* 2013; Ohkuni *et al.* 2016; Ciftci-Yilmaz *et al.* 2018; Au *et al.* 2020; Eisenstatt *et al.* 2020). Growth on galactose media revealed that the SDL of *doa1Δ, slx5Δ, cdc4-1*, and *cdc7-4 GALCSE4* strains is suppressed when either *HHF1* or *HHF2* is deleted (Figures 2E and 2F and Table 2), while the SDL of *hir2Δ GALCSE4* is suppressed only when *HHF2* is deleted (Figure 2E and Table 2). These results suggest that the gene dosage of *H4* contributes to the SDL of mutants that exhibit defects in Cse4 proteolysis and mislocalize Cse4 to non-centromeric regions.

### Reduced gene dosage of *H4* reduces the mislocalization of Cse4 in *psh1Δ* strains

The SDL phenotype of *psh1Δ GALCSE4* strains is correlated with the mislocalization of Cse4 to non-centromeric regions (Hewawasam *et al.* 2010; Ranjitkar *et al.* 2010). We examined if the suppression of SDL in the *psh1Δ hhf1Δ GALCSE4* or *psh1Δ hhf2Δ GALCSE4* strains is due to reduced mislocalization of Cse4. We performed ChIP-qPCR to assay the localization of Cse4 using chromatin from wild type, *psh1Δ, hhf1Δ, hhf2Δ, psh1Δ hhf1Δ*, and *psh1Δ hhf2Δ* strains transiently overexpressing *CSE4*. In agreement with previously published data (Hildebrand and Biggins 2016; Hewawasam *et al.* 2018; Ohkuni *et al.* 2020), we found that Cse4 enrichment at non-centromeric regions such as the promoters of *RDS1, SLP1, GUP2*, and *COQ3* is higher in the *psh1Δ* strain compared to the wild type strain (Figures 3A and 3B and S4A and S4B). In contrast, deletion of *HHF2* in a wild type strain or when combined with *psh1Δ* showed reduced levels of Cse4 enrichment at these regions (Figures 3A and 3B). Results for ChIP-qPCR with the *hhf1Δ* strain also showed reduced levels of Cse4 at non-centromeric loci (Figure S4A and S4B). Consistent with previous studies (Hildebrand and Biggins 2016), we observed higher levels of Cse4 at peri-centromeric regions in a *psh1Δ* strain (Figures 3C and S4C). However, we observed reduced levels of Cse4 at peri-centromeric regions in *psh1Δ hhf1Δ* and *psh1Δ hhf2Δ* strains when compared to the *psh1Δ* strain (Figures 3C and S4C). Localization of Cse4 to the centromere was not significantly altered in *hhf1Δ, hhf2Δ, psh1Δ hhf1Δ*, and *psh1Δ hhf2Δ* strains (Figures 3C and S4C). Based on these results, we conclude that reduced gene dosage of *H4* contributes to reduced levels of Cse4 at non-centromeric and peri-centromeric regions in *psh1Δ* strains.

**Figure 3.**
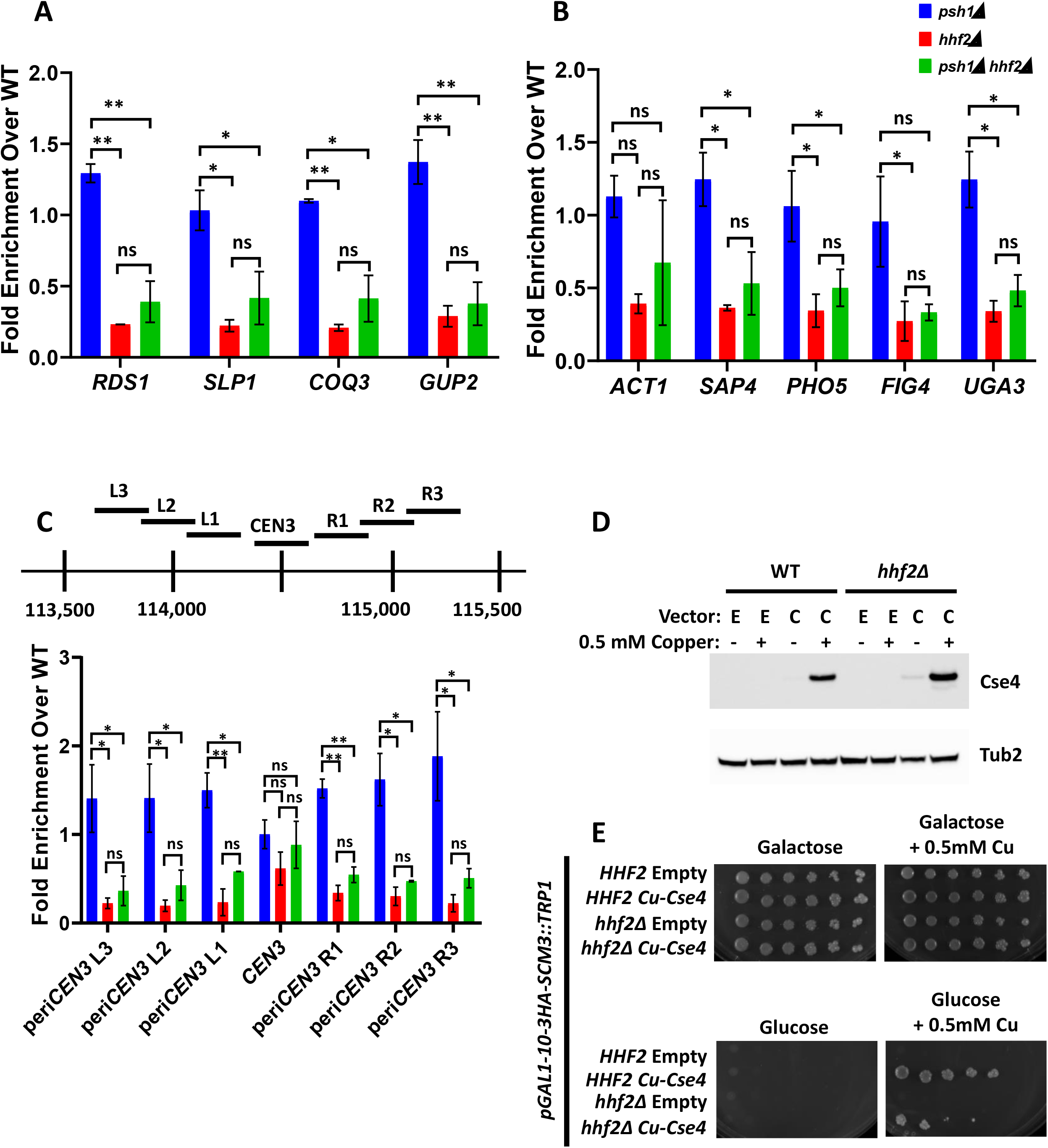
Deletion of *HHF2* reduces enrichment of Cse4 at peri-centromeric and non-centromeric regions. (A-C) ChIP-qPCR was performed on chromatin lysate from wild type (YMB9804), *psh1Δ* (YMB10479), *hhf2Δ* (YMB10938), and *psh1Δ hhf2Δ* (YMB10824) strains transiently overexpressing *GAL1-6His-3HA-CSE4* (pMB1458). Enrichment of 6His-3HA-Cse4 is shown as a fold over wild type. Displayed are the mean of two independent experiments. Error bars represent standard deviation of the mean. \*\**p*-value < 0.0099, \**p*-value < 0.09, ns=not significant. **A.** and **B. Levels of Cse4 enrichment at non-centromeric regions are reduced in a *hhf2Δ* strain.** Enrichment of 6His-3HA-Cse4 at (A) *RDS1, SLP1, COQ3, GUP2*, and (B) *ACT1, SAP1, PHO5, FIG4*, and *UGA3*. **C. Levels of Cse4 at peri-centromeric regions, but not at the core centromere, are significantly reduced when *HHF2* is deleted.** Top: A diagram of the peri-centromere and centromere of Chromosome III analyzed by ChIP-qPCR. Horizontal lines represent the regions amplified. Bottom: Enrichment of 6His-3HA-Cse4 at the core centromere and at the left and right peri-centromeric regions on Chromosome III. **D. Cse4 is expressed from a copper-inducible promoter in *hhf2Δ* strains depleted of Scm3.** Strains from (D) were grown to logarithmic phase in liquid media selective for the plasmid. Cells were induced with 0.5 mM copper for 2 hours and protein lysates were collected and analyzed by Western blot against Cse4 and Tub2 as a loading control. E: empty vector; C: copper inducible Cse4; -: no copper; +: 0.5 mM copper. **E. Deletion of *HHF2* reduces Cse4 deposition at the centromere in cells depleted of Scm3.** Growth assays of strains in which Scm3 is expressed from a galactose inducible promoter and Cse4 is expressed from a copper-inducible promoter. Wild type and *hhf2Δ* with empty vector (pSB17; JG1589 and YMB11252, respectively) or a plasmid with copper inducible Cse4 (pSB873; JG1690 and YMB11254, respectively) were plated in five-fold serial dilutions on media plates selective for the plasmid with raffinose/galactose (2% final concentration each) or glucose (2 % final concentration) and with or without copper (0.5 mM final concentration). Plates were grown for three to five days at 30°C. Two independent transformants were tested and a representative image is shown.

Scm3 is the primary chaperone for centromeric deposition of Cse4 and strains depleted for Scm3 are not viable (Camahort *et al.* 2007). However, overexpression of Cse4 can rescue the growth defect of Scm3-depleted cells, suggesting that non-Scm3-based mechanisms can promote centromeric deposition of overexpressed Cse4 (Hewawasam *et al.* 2018). Our studies so far have shown that reduced gene dosage of *H4* contributes to suppression of Cse4 mislocalization to non-centromeric regions. We next asked if the reduced gene dosage of *H4* would affect the Scm3-independent centromeric deposition of Cse4 by assaying the growth of Scm3-depleted cells that overexpress *CSE4*. In these strains, expression of Scm3 is regulated by a galactose-inducible promoter and is only expressed when grown in galactose medium, but not in glucose medium. However, overexpression of Cse4 from a copper-inducible promoter can suppress the growth defect caused by depletion of Scm3 on copper-containing medium (Hewawasam *et al.* 2018). We constructed *hhf2Δ GAL-SCM3 Cu-CSE4* strains and performed Western blot analysis to confirm the induced overexpression of Cse4 in these strains when grown in copper-containing medium (Figure 3D). Growth assays showed that deletion of *HHF2* resulted in poor growth of cells when Cse4 is overexpressed in Scm3-depleted strains (Figure 3E, glucose + 0.5mM Cu). We conclude that physiological levels of histone H4 are required for centromeric association of Cse4 in cells depleted of Scm3 and for mislocalization of Cse4 to peri-centromeric and non-centromeric regions in *psh1Δ* strains.

### Deletion of *HHF2* contributes to reduced stability of Cse4 in a *psh1Δ* strain

The SDL phenotype and mislocalization of Cse4 in a *psh1Δ GALCSE4* strain is associated with a higher stability of Cse4 (Hewawasam *et al.* 2010; Ranjitkar *et al.* 2010). The suppression of the *psh1Δ GALCSE4* SDL and the reduced mislocalization of Cse4 by *hhf2Δ* led us to hypothesize that the stability of Cse4 would be reduced in a *psh1Δ hhf2Δ* strain. Protein stability assays showed that, in agreement with previous studies (Hewawasam *et al.* 2010; Ranjitkar *et al.* 2010), transiently overexpressed Cse4 is highly stable in the *psh1Δ* strain when compared to that observed in a wild type strain. The stability of Cse4 was not significantly affected in the *hhf2Δ* strain when compared to the wild type strain. Consistent with our hypothesis, we observed reduced stability of Cse4 in the *psh1Δ hhf2Δ* strain compared to the *psh1Δ* strain (Figure 4A). These results show a correlation between suppression of SDL of a *psh1Δ GALCSE4* strain, lower levels of mislocalized Cse4 at non-centromeric regions, and reduced stability of Cse4 due to reduced gene dosage of *H4*.

**Figure 4.**
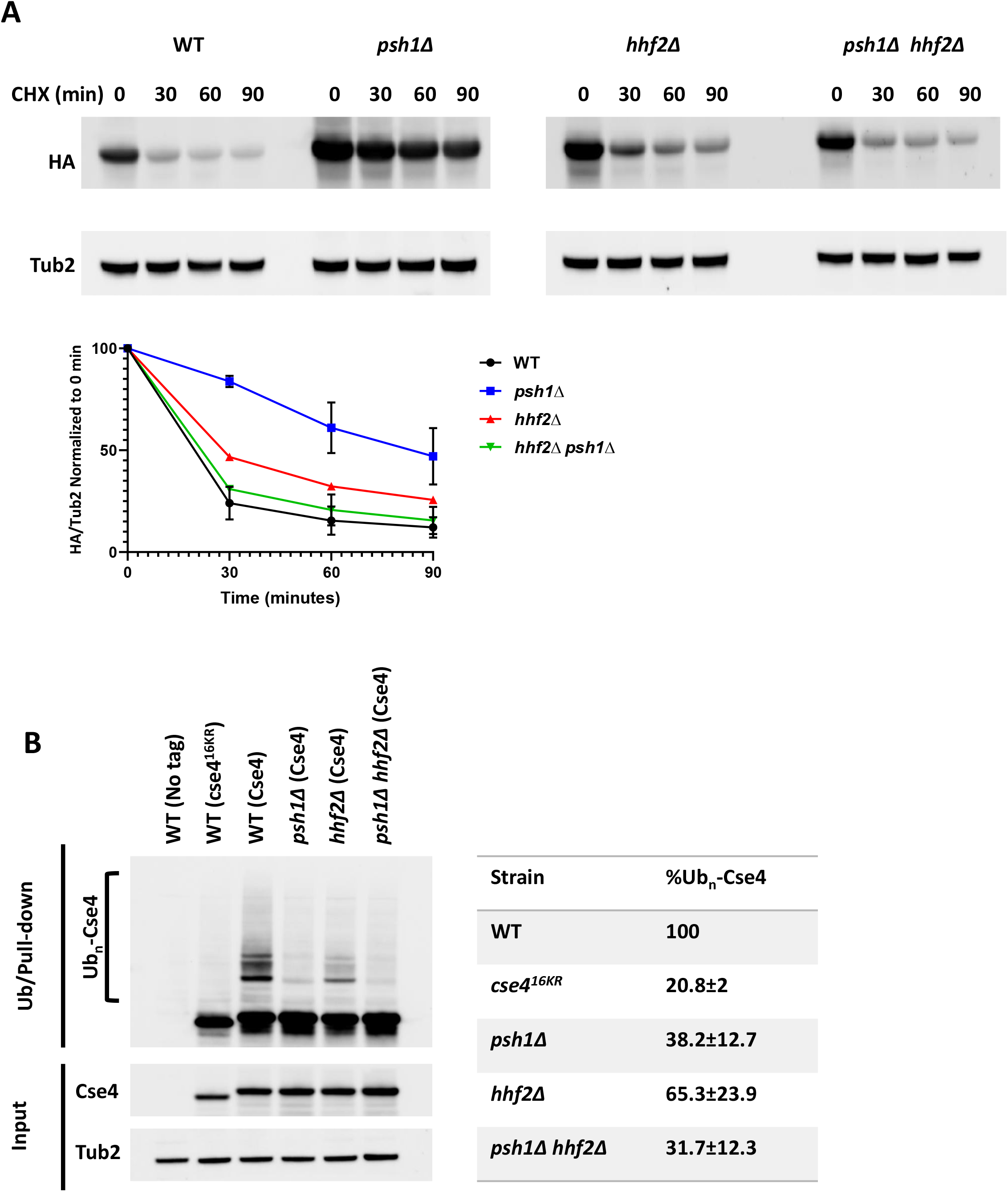
Deletion of *HHF2* contributes to reduced stability and ubiquitin-independent proteolysis of Cse4 in a *psh1Δ* strain. **A. *hhf2Δ* strains contribute to reduced stability of Cse4 in a *psh1Δ* strain.** Western blot analysis of protein extracts from wild type (YMB9804), *psh1Δ* (YMB10479), *hhf2Δ* (YMB10938), and *psh1Δ hhf2Δ* (YMB10824) strains transiently overexpressing *GAL1-6His-3HA-CSE4* (pMB1458). Cells were grown to logarithmic phase in media selective for the plasmid and containing raffinose (2% final concentration) and induced with galactose (2% final concentration) for 4 hours. Cultures were treated with cycloheximide (CHX, 10 µg/mL) and glucose (2%) and analyzed at the indicated time points. Extracts were analyzed by Western blot against HA (Cse4) and Tub2 as a loading control. Levels of 6His-3HA-Cse4 were normalized to Tub2 and the quantification of the percent remaining 6His-3HA-Cse4 after CHX treatment is shown in the graph. Error bars represent the SEM of two independent experiments. **B. Deletion of *HHF2* does not increase ubiquitination of Cse4 in a *psh1Δ* strain.** Ubiquitin-pull down assays were performed using protein extracts from wild type strains (BY4741) with no tag (pMB433) or overexpressing *cse4*^*16KR*^ (pMB1892) and from wild type (YMB9804), *psh1Δ* (YMB10479), *hhf2Δ* (YMB10938), and *psh1Δ hhf2Δ* (YMB10824) strains overexpressing *6His-3HA-CSE4* (pMB1458). Lysates were incubated with Tandem Ubiquitin Binding Entity beads (LifeSensors) prior to analysis of ubiquitin-enriched samples by Western blot against HA and input samples against HA and Tub2 as a loading control. Poly-ubiquitinated Cse4 (Ub_n_-Cse4) is indicated by the bracket. HA levels in input samples were normalized to Tub2 levels and quantification of levels of Ub_n_-Cse4 were normalized to the levels of Cse4 in the input. The percentage of Ub_n_-Cse4 from two independent experiments with standard error is shown.

Since defects in the ubiquitin-proteasome mediated proteolysis of Cse4 contribute to its mislocalization and increased stability (Hewawasam *et al.* 2010; Ranjitkar *et al.* 2010), we investigated if deletion of *HHF2* affects ubiquitination of Cse4 (Ub_n_-Cse4) in a *psh1Δ* strain. Ubiquitin pull-down assays were done to determine the levels of Ub_n_-Cse4 in wild type, *psh1Δ, hhf2*, and *psh1Δ hhf2Δ* strains transiently overexpressing *CSE4*. Wild type strains expressing a non-tagged Cse4 or a mutant form of Cse4 (cse4^16KR^) that cannot be ubiquitinated, where the 16 lysine residues are mutated to arginine, were used as negative controls. As previously reported (Hewawasam *et al.* 2010; Ranjitkar *et al.* 2010), levels of Ub_n_-Cse4 were greatly reduced in the *psh1Δ* strain (38.2%±12.7) when compared to the wild type strain. The levels of Ub_n_-Cse4 in the *psh1Δ hhf2Δ* strain (31.7%±12.3) were similar to the *psh1Δ* strain, however Ub_n_-Cse4 levels were decreased in the *hhf2Δ* strain (65.3%±23.9) compared to the levels in the wild type strain (Figure 4B). We propose that reduced mislocalization of Cse4 and ubiquitin-independent proteolysis of Cse4 contribute to reduced stability of Cse4 in a *psh1Δ hhf2Δ GALCSE4* strain.

### Reduced dosage of *H4* is associated with defects in sumoylation of Cse4

We recently reported that Cse4 is sumoylated and that the sumoylation status of Cse4 at residues K215/216 correlates with the SDL of *psh1*Δ *GALCSE4* strains (Ohkuni *et al.* 2020). Overexpression of the sumoylation-defective *cse4K215/216R/A* does not cause SDL in *psh1*Δ, *slx5*Δ, or *hir2*Δ strains; the lack of an SDL phenotype in the *psh1*Δ strain is due to reduced mislocalization and lower protein stability of cse4K215/216R/A. The phenotypic consequences related to defects in Cse4 sumoylation are similar to the ones we have observed due to reduced dosage of *H4.* We examined if sumoylation of Cse4 is affected due to reduced dosage of *H4*. Wild type, *hhf1Δ*, and *hhf2Δ GALCSE4* strains were assayed for Cse4 sumoylation. Consistent with previous results (Ohkuni *et al.* 2016; Ohkuni *et al.* 2018; Ohkuni *et al.* 2020), we detected sumoylated Cse4 as a pattern of three high molecular weight bands in wild type cells overexpressing wild type Cse4 but not in wild type cells expressing vector alone or overexpressing *cse4*^*16KR*^ (Figure 5A). Deletion of either histone H4 allele resulted in reduced levels of sumoylated Cse4 (Figures 5A and 5B; *p*-value WT vs *hhf1Δ* = 0.0006, *p*-value WT vs *hhf2Δ* = 0.0007). To confirm that the reduction of sumoylated Cse4 is linked to deletion of the histone H4 allele, we assayed the levels of sumoylated Cse4 in *hhf2Δ GALCSE4* strains transformed with an empty vector or with a plasmid borne *HHF2*. As expected, plasmid borne *HHF2* restored the levels of sumoylated Cse4 to that observed in wild type cells (Figures 5C and 5D). We conclude that physiological levels of histone H4 are required for Cse4 sumoylation.

**Figure 5.**
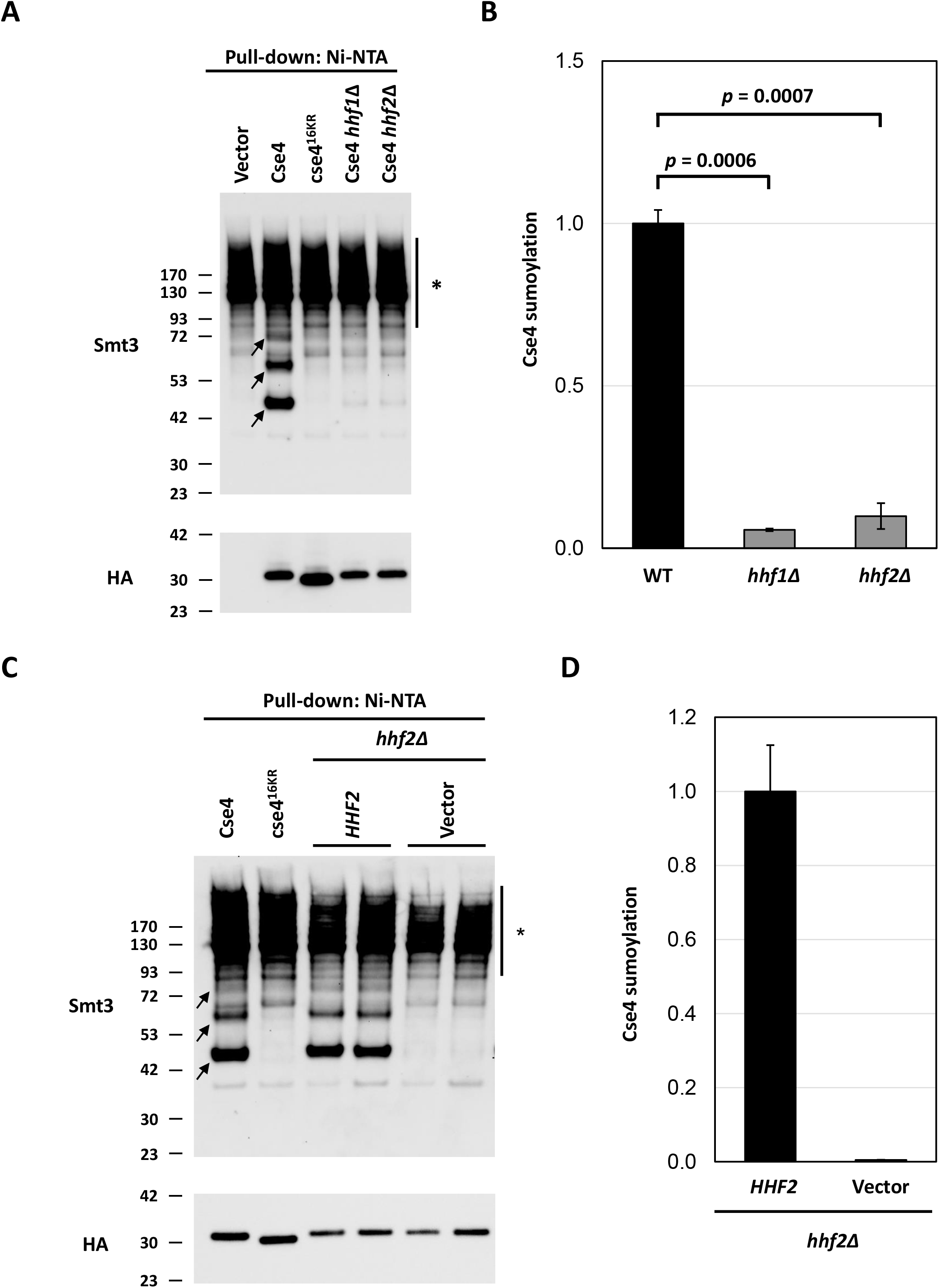
Histone H4 contributes to the sumoylation of Cse4. **A. Levels of sumoylated Cse4 are decreased in histone H4 deletion strains.** Sumoylation levels were assayed on wild type (BY4741) strains transformed with empty vector (pYES2), *pGAL-8His-HA-CSE4* (pMB1345), or *pGAL-8His-HA-cse4*^*16KR*^ (pMB1344) and *hhf1Δ* (YMB10766) and *hhf2Δ* (YMB10767) strains transformed with *pGAL-8His-HA-Cse4* (pMB1345). Sumoylated and nonmodified Cse4 were detected using cell lysates that were incubated with Ni-NTA beads followed by western blot analysis with antibodies against Smt3 and HA (Cse4), respectively. Arrows indicate the three high molecular weight bands that represent sumoylated Cse4. Asterisk indicates nonspecific sumoylated proteins that bind to beads. **B. Quantification of relative levels of sumoylated Cse4 in histone H4 deletion strains.** Levels of sumoylated Cse4 were normalized to non-modified Cse4 probed against HA in the pull-down sample. Statistical significance from two independent experiments was assessed by one-way ANOVA (*p* value = 0.0004) followed by Tukey post test (all pairwise comparisons of means). Error bars indicate average deviation from the mean. **C. The Cse4 sumoylation defect in a *hhf2Δ* strain is linked to the *HHF2* allele.** Sumoylation levels were determined from lysates from a *hhf2*Δ (YMB10767) strain with *pGAL-8His-HA-CSE4* (pMB1345) transformed with vector (pRS425) or *HHF2* (pMB1929) as described in (A). Arrows indicate the three high molecular weight bands that represent sumoylated Cse4. Asterisk indicates nonspecific sumoylated proteins that bind to beads. **D. Quantification of relative levels of sumoylated Cse4.** Relative levels of sumoylated Cse4 were normalized to non-modified Cse4 probed against HA in the pull-down sample. Error bars indicate average deviation from the mean from two biological replicates.

### A histone H4 mutant defective for interaction with Cse4 suppresses the *psh1Δ* ***GALCSE4* SDL and shows defects in Cse4 sumoylation**

Our results so far have shown that reduced gene dosage of *H4* contributes to the suppression of the SDL phenotype, reduced stability of Cse4, decreased mislocalization of Cse4 in *psh1Δ GALCSE4* strains, and defects in Cse4 sumoylation. We hypothesized that strains with defects in the interaction of H4 with Cse4 will display the same phenotypes that are observed due to reduced dosage of *H4* in *psh1*Δ strains. To test our hypothesis, we used *HHT1/hhf1 hht2*Δ*/hhf2*Δ strains with mutations either in the N-terminal lysines (*HHT1/hhf1-10*) or in the histone fold domain (*HHT1/hhf1-20*) (Figure 6A) that have been well characterized by genetic and biochemical analysis (Smith *et al.* 1996; Glowczewski *et al.* 2000). The temperature sensitivity of the *HHT1/hhf1-20* strain, but not the *HHT1/hhf1-10* strain, is suppressed by overexpression of Cse4 and the *HHT1/hhf1-20* strain is proposed to have defects in the formation of the Cse4-H4 dimer (Smith *et al.* 1996; Glowczewski *et al.* 2000). We deleted *PSH1* in the same genetic background as the *HHT1/HHF1, HHT1/hhf1-10*, and *HHT1/hhf1-20* strains, transformed these strains with *CSE4* on a galactose-inducible plasmid, and performed growth assays. Compared to wild type strains with a single copy of genes encoding histones H3/H4, *HHT1/HHF1 psh1Δ* strains display SDL on galactose medium when Cse4 is overexpressed, though to a less prominent degree compared to strains expressing both alleles encoding H3/H4 (compare Figure 6B to Figure 2A, *psh1Δ GALCSE4*). The relative decrease in SDL may be due to the expression of a single copy of the genes encoding histones H3/H4 in the strain background. The *HHT1/hhf1-20* mutant suppresses the SDL of *psh1Δ GALCSE4* strains while the *HHT1/hhf1-10* mutant does not (Figure 6B). These findings suggest that the defect in the Cse4-H4 interaction contributes to the suppression of the *psh1Δ GALCSE4* SDL in the *HHT1/hhf1-20* strain.

**Figure 6.**
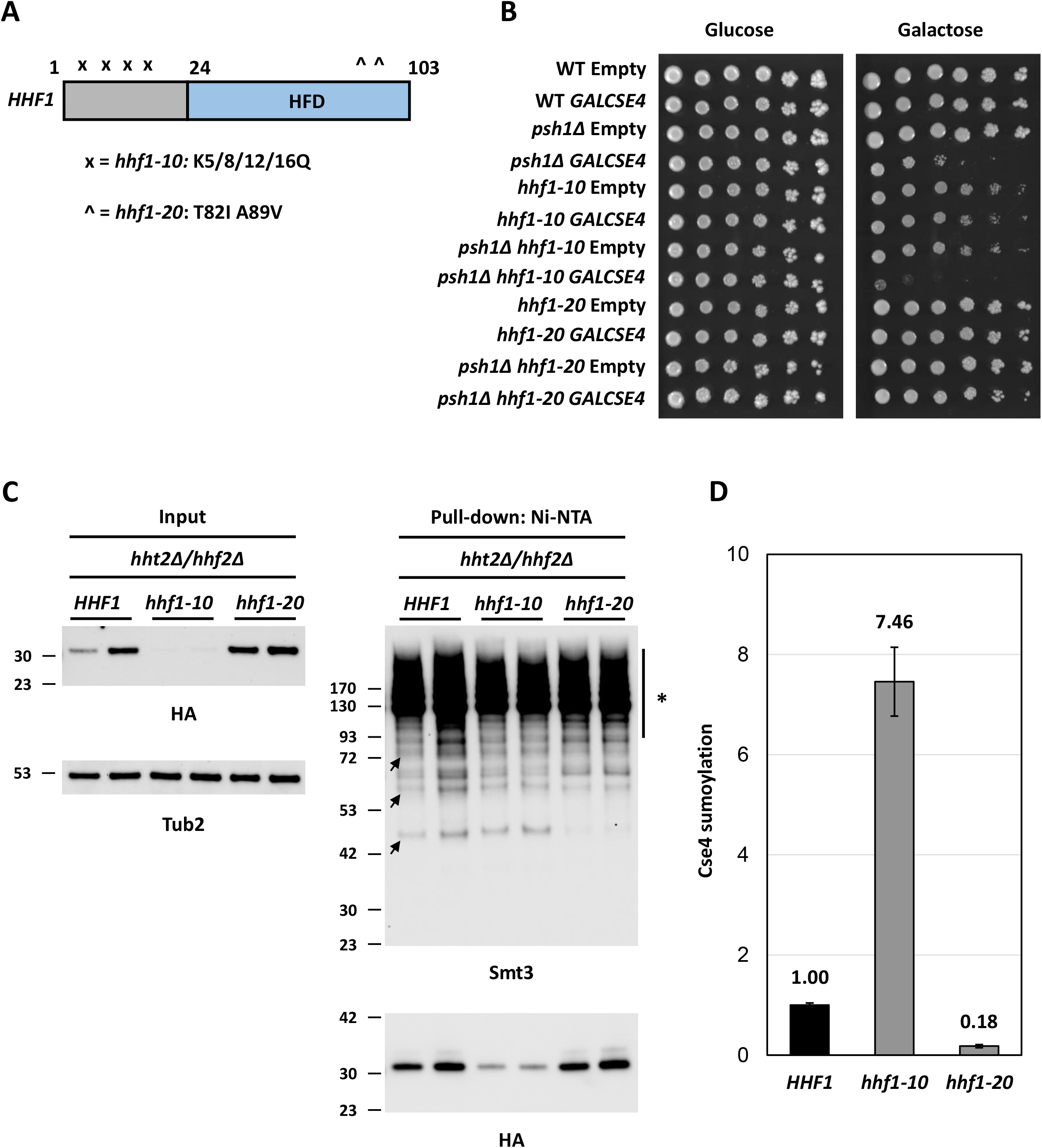
Mutation in the histone fold domain of H4 histone suppresses the SDL phenotype of a *psh1*Δ *GALCSE4* strain and causes defects in Cse4 sumoylation. A. Schematic of *HHF1*. Displayed is a cartoon of the *HHF1* gene with mutations in *hhf1-10* indicated by an ‘x’ and *hhf1-20* with a ‘^’ in the histone fold domain (HFD, blue). The specific residues mutated in each allele are indicated below the schematic. **B. Mutations in the histone fold domain of histone H4 suppress the SDL phenotype of a *psh1Δ GALCSE4* strain.** Growth assays of wild type (MSY559), *psh1Δ* (YMB11346), *HHT1/hhf1-10* (MSY535), *HHT1/hhf1-20* (MSY534), *psh1Δ HHT1/hhf1-10* (YMB11347), and *psh1Δ HHT1/hhf1-20* (YMB11348) with empty vector (pMB433) or expressing *GAL1-6His-3HA-CSE4* (pMB1458). Cells were plated in five-fold serial dilutions on selective media plates containing either glucose (2% final concentration) or raffinose/galactose (2% final concentration each). Plates were incubated at 30°C for three to five days. Three independent transformants were tested and a representative image is shown. **C. Mutations in the histone fold domain of histone H4 decrease levels of sumoylated Cse4.** The levels of sumoylated Cse4 were determined using lysates from *HHT1/HHF1* (MSY559), *HHT1/hhf1-10* (MSY535), and *HHT1/hhf1-20* (MSY534) strains in the *hht2*Δ/*hhf2*Δ background, transformed with *pGAL-8His-HA-CSE4* (pMB1345), as described in Figure 5A. Arrows indicate the three high molecular weight bands that represent sumoylated Cse4. Asterisk indicates nonspecific sumoylated proteins that bind to beads. **D. Quantification of the relative levels of sumoylated Cse4 in *hhf1* strains.** Levels of sumoylated Cse4 were normalized to non-modified Cse4 probed against HA in the pull-down samples and levels in the *HHT1/HHF1* strain were set to 1. Error bars indicate average deviation from the mean from two biological replicates.

We next examined the stability of Cse4 in *HHT1/HHF1, HHT1/HHF1 psh1Δ, HHT1/hhf1-10, HHT1/hhf1-20, HHT1/hhf1-10 psh1Δ*, and *HHT1/hhf1-20 psh1Δ* strains transiently overexpressing *CSE4.* In agreement with previous findings (Figure 4A), overexpressed Cse4 is rapidly degraded in *HHT1/HHF1* cells and is stabilized in the *HHT1/HHF1 psh1Δ* strain (Hewawasam *et al.* 2010; Ranjitkar *et al.* 2010) (Figure S5, top panels). Interestingly, degradation of overexpressed Cse4 in both *HHT1/hhf1-10* and *HHT1/hhf1-20* strains was faster compared to the *HHT1/HHF1* strain. The *HHT1/hhf1-20 psh1Δ* strain showed rapid degradation of Cse4 when compared to the *HHT1/HHF1 psh1Δ* and *HHT1/hhf1-10 psh1*Δ strains (Figure S5). The rapid degradation of overexpressed Cse4 in the *HHT1/hhf1-20 psh1Δ* strain is consistent with previous studies for a correlation between higher protein stability and *GALCSE4* SDL (Hewawasam *et al.* 2010; Ranjitkar *et al.* 2010; Ciftci-Yilmaz *et al.* 2018; Au *et al.* 2020; Eisenstatt *et al.* 2020) and suggests that defective Cse4-H4 interaction contributes to the lack of *GALCSE4* SDL in *psh1*Δ strains.

To examine the effect of the *HHT1/hhf1-20* and *HHT1/hhf1-10* alleles on the levels of Cse4 sumoylation, we used *HHT1/HHF1, HHT1/hhf1-10*, and *HHT1/hhf1-20* strains overexpressing *CSE4* to examine the sumoylation status of Cse4. Western blot analysis was performed after equal amounts of protein (5 mg) for each strain were pulled down with Ni-NTA agarose beads and normalized to the levels of non-modified Cse4 in the pull down (Figure 6C and D). Sumoylated Cse4 was observed in the *HHT1/HHF1* and the *HHT1/hhf1-10* strains (Figures 6C and 6D). Levels of sumoylated Cse4 were normalized to non-modified Cse4 in the pull down samples. The low expression of Cse4 in the *HHT1/hhf1-10* strain (Figure 6C, input) contributes to the higher levels of Cse4 sumoylation due to normalization to the low levels of non-modified Cse4 in this strain (Figure 6D). In contrast, the levels of Cse4 sumoylation were barely detectable in the *HHT1/hhf1-20* strain when compared to the *HHT1/HHF1* strain (Figures 6C and 6D). The reduced sumoylation of Cse4 in the *HHT1/hhf1-20* strain is consistent with the rescue of SDL in the *HHT1/hhf1-20 psh1*Δ *GALCSE4* strain. We conclude that defects in the interaction of hhf1-20 with Cse4 contributes to reduced Cse4 sumoylation and suppression of *psh1*Δ *GALCSE4* SDL due to rapid degradation of Cse4.

### Cse4 mutants defective in the Cse4-H4 interaction do not cause SDL in a *psh1Δ* strain and exhibit defects in Cse4 sumoylation

To further confirm that the Cse4-H4 interaction contributes to SDL in a *psh1*Δ *GALCSE4* strain and Cse4 sumoylation, we investigated if Cse4 residues that are essential for the Cse4-H4 dimer formation (Figure 7A) affect the SDL of a *psh1*Δ strain and sumoylation of Cse4. Like the *HHT1/hhf1-20* mutant, the *cse4* mutants *cse4-102* (L176S M218T) and *cse4-111* (L194Q) exhibit defects in the Cse4-H4 dimer formation, while *cse4-110* (L197S) likely impairs formation of the (Cse4-H4)_2_ tetramer (Glowczewski *et al.* 2000). We hypothesized that overexpression of these *cse4* mutants will not lead to SDL in a *psh1*Δ strain and these mutants will show defects in Cse4 sumoylation. To test these hypotheses, we generated galactose-inducible plasmids expressing *cse4-102* (L176S M218T), *cse4-107*^*MB*^ (L176S), *cse4-108* (M218T), *cse4-110* (L197S), and *cse4-111* (L194Q). To test the effect of the *cse4* mutants on SDL in a *psh1Δ* strain, we performed growth assays. We first determined that overexpression of mutant *cse4* from these plasmids did not result in growth defects in a wild type strain (Figure S6). In agreement with our hypothesis, overexpression of all *cse4* mutants did not cause SDL in a *psh1Δ* strain (Figure 7B). We conclude that the Cse4-H4 dimerization is essential for the SDL phenotype of a *psh1*Δ *GALCSE4* strain.

**Figure 7.**
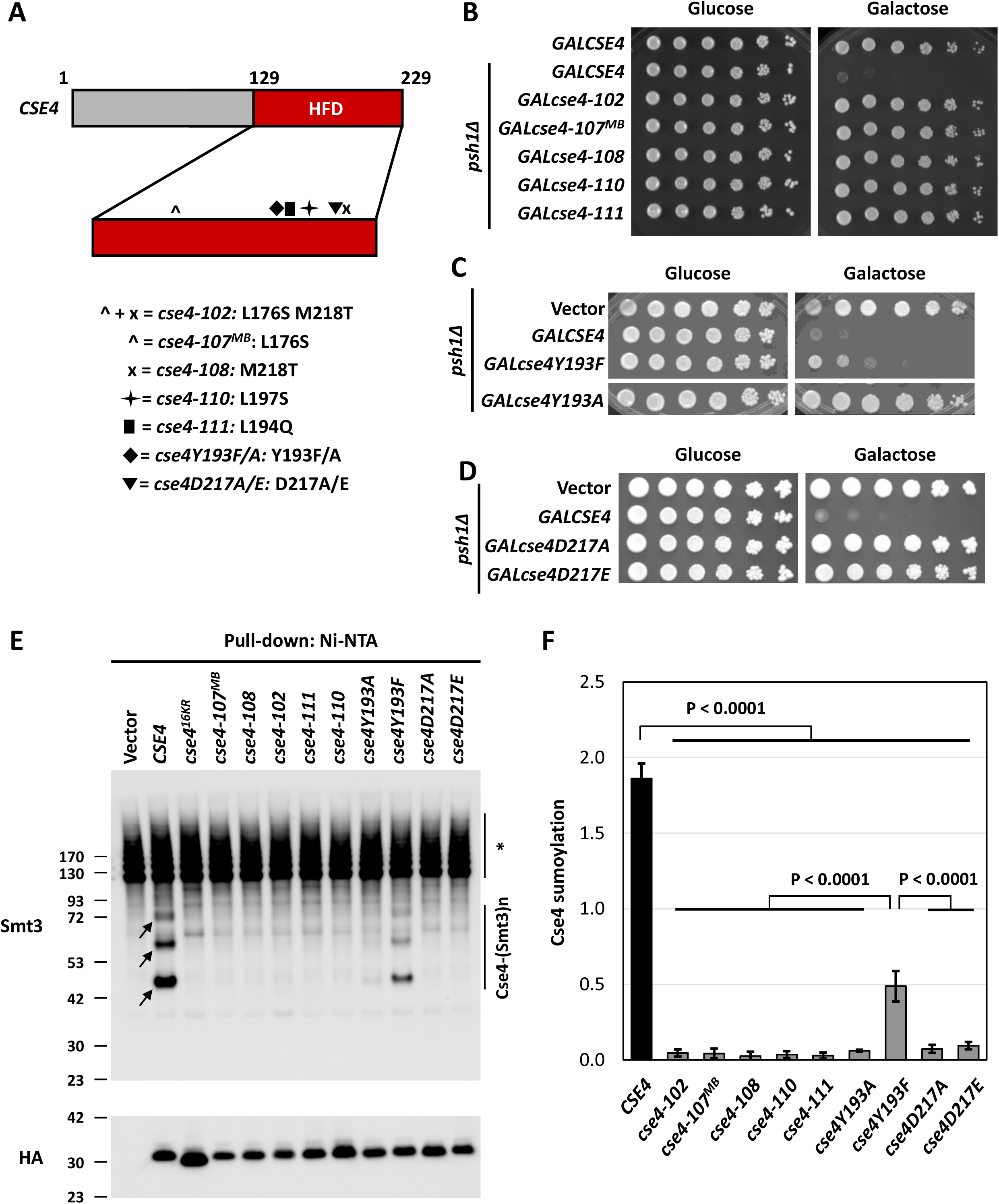
Cse4 mutants defective in the Cse4-H4 interaction do not cause SDL in a *psh1*Δ *GALCSE4* strain and exhibit defects in Cse4 sumoylation. **A. Schematic of *CSE4*.** Displayed is a cartoon of the *CSE4* gene highlighting mutations in the histone fold domain (HFD, red). The HFD is expanded under the representation of *CSE4*. Below the gene schematic is a key describing the symbol that represents a specific mutant *cse4* allele and the residues mutated. **B. Cse4-H4 assembly mutants in Cse4 do not cause SDL in a *psh1Δ GALCSE4* strain.** Growth assays of a *psh1Δ* (YMB8995) strain transformed with *pGAL1-8His-HA-Cse4* (pMB1344), *pGAL1-8His-HA-cse4-102* (pMB1984), *pGAL1-8His-HA-cse4-107*^*MB*^ (pMB1985), *pGAL1-8His-HA-cse4-108* (pMB1986), *pGAL1-8His-HA-cse4-110* (pMB1987), or *pGAL1-8His-HA-cse4-111* (pMB1988). Cells were plated in five-fold serial dilutions on selective media plates containing either glucose (2% final concentration) or raffinose/galactose (2% final concentration each). Plates were incubated at 30°C for three to five days. Three independent transformants were tested and a representative image is shown. **C. The Y193A mutation in Cse4 does not cause SDL in a *psh1Δ GALCSE4* strain.** Growth assays of a *psh1Δ* (YMB9034) strain transformed with empty vector (pYES2), *pGAL1-8His-HA-cse4*^*Y193A*^ (pMB1766), or *pGAL1-8His-HA-cse4Y193F* (pMB1787). Five-fold serial dilutions of the indicated strains were plated on glucose (2% final concentration)- or galactose (2% final concentration)-containing medium selective for the plasmid. The plates were incubated at 30°C for 3days. **D. The *cse4D217A*/E mutants do not cause SDL in a *psh1Δ* strain.** Growth assays of a *psh1Δ* (YMB9034) strain transformed with empty vector (pYES2), *pGAL1-8His-HA-cse4D217A* (pMB1910), or *pGAL1-8His-HA-cse4D217E* (pMB1920). Strains were assayed as described in (C). **E. Cse4 sumoylation levels are decreased in Cse4-H4 assembly mutants.** Levels of sumoylated Cse4 were assayed in a wild type strain (BY4741) transformed with empty vector (pYES2), *pGAL1-8His-HA-CSE4* (pMB1345), *pGAL1-8His-HA-cse4*^*16KR*^ (pMB1344), *pGAL1-8His-HA-cse4-107*^*MB*^ (pMB1985), *pGAL1-8His-HA-cse4-108* (pMB1986), *pGAL1-8His-HA-cse4-102* (pMB1984), *pGAL1-8His-HA-cse4-111* (pMB1988), *pGAL1-8His-HA-cse4-110* (pMB1987), *pGAL1-8His-HA-cse4Y193A* (pMB1766), *pGAL1-8His-HA-cse4Y193F* (pMB1787), *pGAL1-8His-HA-cse4D217A* (pMB1910), or *pGAL1-8His-HA-cse4D217E* (pMB1920). Arrows indicate the three high molecular weight bands that represent sumoylated Cse4. Asterisk indicates nonspecific sumoylated proteins that bind to beads. **F. Quantification of the relative levels of sumoylated Cse4 in *cse4* mutants.** Levels of sumoylated Cse4 in arbitrary density units were normalized to non-modified Cse4 probed against HA in the pull-down samples. Statistical significance from at least three biological repeats was assessed by one-way ANOVA (*p*-value < 0.0001) followed by Tukey post-test (all pairwise comparisons of means). Error bars indicate standard deviation from the mean.

Next, we generated galactose-inducible plasmids expressing *cse4Y193A/F* and *cse4D217A/E* for growth assays in a *psh1*Δ strain. The rationale for *cse4Y193A/F* is that Y193 is next to the mutated residue in *cse4-111* (L194Q), is located at the center of the α2 helix of Cse4, and interacts with the α2 helix of H4 in the context of Scm3 (Zhou *et al.* 2011). For *cse4D217A/E*, the D217 residue is adjacent to the residue mutated in *cse4-108* (M218T), is part of the K215/216 sumoylation consensus site, 214-M**KK**D-217 (Ψ-K-x-D/E), and is essential for dimerization of Cse4 (Camahort *et al.* 2009). Growth assays on galactose media showed that *cse4Y193A* and *cse4D217A/E* do not cause SDL in a *psh1*Δ strain (Figures 7C and 7D). Note that *cse4Y193F* showed partial lethality in a *psh1*Δ strain when compared to *CSE4* (Figure 7C). Taken together, these results show that overexpression of the *cse4* mutants with defects in the formation of the Cse4-H4 dimer, do not lead to a SDL phenotype in a *psh1*Δ strain.

The lack of SDL in *psh1Δ* strains overexpressing *cse4-102, cse4-107*^*MB*^, *cse4-108, cse4-110, cse4-111, cse4Y193A*, and *cse4D217A/E* is similar to the suppression of *psh1Δ GALCSE4* SDL when combined with *hhf1*Δ, *hhf2*Δ, and *hhf1-20 s*trains. Defects in Cse4 sumoylation in *hhf1Δ, hhf2Δ*, and *hhf1-20* strains led us to hypothesize that *cse4-102, cse4-107*^*MB*^, *cse4-108, cse4-110, cse4-111, cse4Y193A*, and *cse4D217A/E* strains will also show defects in Cse4 sumoylation. Thus, we examined the sumoylation status of the *cse4* mutants used in the growth assays (Figure 7E). Consistent with our hypothesis, levels of Cse4 sumoylation were reduced in all *cse4* mutants except *cse4Y193F*, which showed only a partial reduction of Cse4 sumoylation (Figures 7E and 7F). The reduced sumoylation of cse4Y193F is consistent with the partial lethality observed in a *psh1Δ* strain expressing *cse4Y193F*. Our results demonstrate that overexpression of *cse4* mutants defective for the Cse4-H4 dimer formation lead to defects in Cse4 sumoylation. We conclude that the Cse4-H4 dimer formation regulates Cse4 sumoylation and this contributes to *psh1*Δ *GALCSE4* SDL.

## DISCUSSION

Mislocalization of overexpressed CENP-A and its homologs contributes to chromosomal instability (CIN) in yeast, fly, and human cells (Heun *et al.* 2006; Au *et al.* 2008; Mishra *et al.* 2011; Lacoste *et al.* 2014; Athwal *et al.* 2015; Shrestha *et al.* 2017) and overexpression and mislocalization of CENP-A are observed in many cancers (Tomonaga *et al.* 2003; Amato *et al.* 2009; Li *et al.* 2011; McGovern *et al.* 2012; Sun *et al.* 2016; Zhang *et al.* 2016). In this study, we performed the first genome-wide screen to identify deletion or temperature sensitive (*ts*) mutants that suppress the synthetic dosage lethality (SDL) due to mislocalization of overexpressed Cse4 in *psh1*Δ *GALCSE4* strains. Deletion of either allele that encodes histone H4 (*HHF1* and *HHF2*) were among the most prominent suppressors of *psh1Δ GALCSE4* SDL. We determined that reduced gene dosage of *H4* contributes to defects in Cse4 sumoylation and this prevents mislocalization of overexpressed Cse4 at peri-centromeric and non-centromeric regions, leading to suppression of the *psh1Δ GALCSE4* SDL. We also determined that the Cse4-H4 interaction contributes to Cse4 sumoylation and *psh1Δ GALCSE4* SDL as *hhf1-20, cse4-102*, and *cse4-111* mutants, which are defective for the Cse4-H4 interaction, exhibit reduced sumoylation of Cse4 and do not exhibit *psh1Δ GALCSE4* SDL. Taken together, our genome-wide screen identified genes that contribute to Cse4 mislocalization and provides mechanistic insights into how reduced gene dosage of *H4* prevents mislocalization of Cse4 into non-centromeric regions.

The suppressor screen was performed under a condition with high levels of Cse4 expression induced from a *GAL1-6His-3HA-CSE4* plasmid, which contributes to mild growth sensitivity even in wild type cells and this leads to lethality in *psh1Δ* strains (Figure 2). To reduce the number of false positive suppressors, we performed the screen with a *psh1Δ GALCSE4* strain grown on 2% galactose medium to achieve maximum levels of Cse4 overexpression. These growth conditions limited us from identifying partial suppressors such as deletion of *NHP10*, which encodes a subunit of the INO80 chromatin remodeling complex and was previously shown to suppress the *psh1Δ GALCSE4* SDL on medium with a lower concentration of galactose (0.1%) (Hildebrand and Biggins 2016). While our screen did not identify *nhp10Δ*, it did identify two deletions and one mutant allele for genes encoding INO80 subunits, Ies2, Arp8, and Act1, respectively, that are evolutionarily conserved between yeast and human cells (Poch and Winsor 1997; Shen *et al.* 2000; Shen *et al.* 2003; Tosi *et al.* 2013). Secondary growth validation showed that *arp8Δ*, but not *act1-132* or *ies2Δ*, suppresses the *psh1Δ GALCSE4* SDL. However, the polyploid nature of the *arp8Δ* strain precluded further study with this suppressor. The stringent growth conditions of the screen also prevented the identification of deletion of Cac2, a subunit of the CAF-1 complex, which promotes Cse4 incorporation at non-centromeric regions (Hewawasam *et al.* 2018). We determined that *cac2Δ* cannot suppress the *psh1Δ GALCSE4* SDL under the conditions used in our screen (data not shown).

Previous studies have shown that mislocalization of Cse4 to non-centromeric regions contributes to the *GALCSE4* SDL in *psh1Δ, slx5Δ, doa1Δ, hir2Δ, cdc4-1*, and *cdc7-4* strains (Hewawasam *et al.* 2010; Ranjitkar *et al.* 2010; Au *et al.* 2013; Ohkuni *et al.* 2016; Ciftci-Yilmaz *et al.* 2018; Au *et al.* 2020; Eisenstatt *et al.* 2020). We sought to define mechanisms that prevent lethality due to mislocalization of overexpressed Cse4. The identification of both *hhf1Δ* and *hhf2Δ* as suppressors of *psh1Δ GALCSE4* SDL led us to examine how reduced gene dosage of *H4* contributes to preventing mislocalization of Cse4. A role for histone H4 in centromeric localization of Cse4 has been examined previously (Deyter *et al.* 2017) however, the effect of gene dosage of *H4* in non-centromeric chromosome localization of Cse4 has not yet been explored. We determined that suppression of the *GALCSE4* SDL phenotype by *hhf1*Δ and *hhf2*Δ is not restricted to *psh1*Δ strains and is also observed in *slx5Δ, doa1Δ, cdc4-1*, and *cdc7-4* strains. The SDL phenotype of the *hir2Δ GALCSE4* strain showed better suppression with *hhf2Δ* than with *hhf1Δ*. This may be due to differential expression of H4 mRNA, which is five to seven times more abundant from the *HHT2-HHF2* allele than from the *HHT1-HHF1* allele (Cross and Smith 1988) or due to the role of the HIR complex in histone gene expression (Prochasson *et al.* 2005; Fillingham *et al.* 2009; Kurat *et al.* 2014).

We used several approaches to understand the molecular mechanism for suppression of the *psh1Δ GALCSE4* SDL phenotype by *hhf1Δ* and *hhf2Δ*. These include ChIP-qPCR at regions of known Cse4 association, protein stability assays, and determining the status of Cse4 ubiquitination and sumoylation. Genome-wide studies have shown that overexpressed Cse4 is significantly enriched at promoters and peri-centromeric regions in a *psh1*Δ strain (Hildebrand and Biggins 2016). Our ChIP-qPCR data showed reduced levels of Cse4 at peri-centromeric and non-centromeric regions in *psh1Δ hhf1Δ* and *psh1Δ hhf2Δ* strains when compared to the *psh1Δ* strain. The mislocalization of overexpressed Cse4 to non-centromeric regions contributes to highly stable Cse4 in *psh1Δ, slx5Δ, doa1Δ, hir2Δ, cdc4-1*, and *cdc7-4* strains (Hewawasam *et al.* 2010; Ranjitkar *et al.* 2010; Au *et al.* 2013; Ohkuni *et al.* 2016; Ciftci-Yilmaz *et al.* 2018; Au *et al.* 2020; Eisenstatt *et al.* 2020). We reasoned that reduced mislocalization of Cse4 to non-centromeric regions in *psh1Δ hhf2*Δ strains may contribute to faster degradation of Cse4 in these strains. Our results showed that the proteolysis of Cse4 was indeed faster in *psh1*Δ *hhf2*Δ strains when compared to the *psh1*Δ strain. Intriguingly, this was not due to increased ubiquitination of Cse4 (Ub_n_-Cse4) in *psh1*Δ *hhf2*Δ strains. These results suggest a ubiquitin-independent mechanism that may contribute to the proteolysis of Cse4 in *hhf2Δ psh1Δ* strains. Ubiquitin-independent proteolysis has also been reported previously as cse4^16KR^, in which all lysine residues are mutated to arginine, is still degraded (Collins *et al.* 2004).

Our results showing that reduced dosage of *H4* contributes to the suppression of *GALCSE4* SDL in *psh1Δ* strains, reduced mislocalization of Cse4, and lower protein stability of Cse4 are similar to the phenotypes of the sumoylation-defective *cse4K215/216R/A* strains (Ohkuni *et al.* 2020). Consistent with these results, deletion of either histone H4 allele resulted in reduced levels of sumoylated Cse4. We therefore propose that physiological levels of H4 regulate the sumoylation of Cse4 and that this in turn facilitates mislocalization of overexpressed Cse4 to non-centromeric regions and *GALCSE4* SDL in mutant such as *psh1Δ*. Importantly, in contrast to histone H4, reduced dosage of genes encoding other canonical histones such as histones H2A or H3 does not suppress the *psh1Δ GALCSE4* SDL.

To further examine the role of *H4* in regulating the mislocalization of Cse4, we pursued studies using well characterized separation of function alleles of *H4 (hhf1-20)* and *CSE4 (cse4-102* and *cse4-111)* with defects in the Cse4-H4 interaction (Smith *et al.* 1996; Glowczewski *et al.* 2000). Consistent with a role of H4 for its interaction with Cse4, we observed suppression of the *psh1Δ GALCSE4* SDL in a *hhf1-20* strain and lack of SDL when *cse4-102* or *cse4-111* were overexpressed in a *psh1Δ GALCSE4* strain. The *hhf1* mutant strains lack the *HHT2/HHF2* allele and express only a single copy of *H3/H4* (*HHT1/HHF1*). In this strain background, the *psh1Δ GALCSE4* SDL was less severe compared to results in our strains with wild type copies of both *HHT1/HHF1* and *HHT2/HHF2* (Figure 2). Despite this, we were able to unambiguously establish that *HHT1/hhf1-20*, but not *HHT1/hhf1-10*, suppresses the *psh1Δ GALCSE4* SDL. Interestingly, the *hhf1-10 psh1Δ GALCSE4* strain displayed a more lethal phenotype than the wild type *HHT1/HHF1 psh1Δ GALCSE4* strain. The N-terminal lysine residues on histone H4 (K5, 8, 12, 16) are acetylated and the *HHT1/hhf1-10* mutations mimic the hyperacetylated state of the lysine residues (K to Q). We have previously shown that levels of acetylated H4 are low at centromeres and that the maintenance of hypoacetylated H4 at the centromere is essential for kinetochore function and faithful chromosome segregation (Choy *et al.* 2011). We propose that the hyperacetylated state of H4 in the *HHT1/hhf1-10* strain contributes to the more severe SDL that we observed. A recent study showed that strains with a mutation of histone H4 arginine 36 to alanine (H4R36A) display SDL when Cse4 is overexpressed and that this is due to defects in the interaction of H4R36A with Psh1, thereby leading to enrichment of Cse4 and Psh1 at non-centromeric regions in these cells (Deyter *et al.* 2017).

Consistent with our previous studies (Ohkuni *et al.* 2020), we observed a correlation between the suppression of *GALCSE4* SDL and reduced sumoylation of Cse4 in *HHT1/hhf1-20, cse4-102*, and *cse4-111* strains. Similar results were observed with *cse4Y193A/F*, which is adjacent to the mutated site in *cse4-111* (L194Q), and with *cse4D217D/E*, which is adjacent to the residue mutated in *cse4-108* (M218T) and a part of the K215/216 sumoylation consensus site (Camahort *et al.* 2009). Accordingly, low levels of sumoylated cse4Y193F correlate with a partial lethality of a *psh1Δ GALcse4Y193F* strain and severe defects in sumoylated cse4Y193A correlate with a lack of SDL in a *psh1Δ GALcse4Y193A* strain. Phenylalanine (F) is identical to tyrosine (Y) except for the hydroxyl group present on Y. It is possible that the structural similarity between Y and F allows at least partial formation of the Cse4-H4 dimer, resulting in partial sumoylation of cse4Y193F. In contrast, we observed a reduction of Cse4 sumoylation of both cse4D217A and cse4D217E mutants compared to wild type. The D217 residue of Cse4 is essential for growth and is important for the Cse4 dimerization. Since the *cse4D217E* mutant, which is part of the intact sumoylation consensus site, shows reduction of Cse4 sumoylation and does not complement the null mutation (Figure S7), we propose that D217 has a role besides regulating sumoylation of Cse4K215/216. Sumoylation of Cse4 is not essential for centromeric localization of Cse4 because a *cse4*^*16KR*^ strain with all 16 lysine (K) residues mutated to arginine (R) is viable in the context of the wild type centromeric chaperone Scm3 (Au *et al.* 2008). Sumoylation of Cse4K215/216 or physiological levels of H4 are indispensable only when Scm3 is not expressed (Ohkuni *et al.* 2020). Our results show that defects in Cse4 sumoylation contribute to reduced levels of non-centromere associated Cse4 with no significant effect on levels of centromere associated Cse4 in *psh1Δ hhf1Δ* and *psh1Δ hhf2Δ* strains. We propose that reduced dosage of *H4* serves to protect the cells from the detrimental effects of overexpressed Cse4 due to defects in Psh1, SCF^Cdc4^, Cdc7, Slx5/8, HIR, and Doa1-mediated proteolysis of Cse4. We define a previously undefined role for histone H4 gene dosage and the Cse4-H4 interaction as key upstream events for the sumoylation of Cse4, which facilitates non-centromeric localization of overexpressed Cse4 and SDL in a *psh1Δ GALCSE4* strain.

In summary, our genome-wide screen identified suppressors of *psh1Δ GALCSE4* SDL with deletions of either allele that encodes histone H4 (*HHF1* and *HHF2)* as among the most prominent suppressors. We present several experimental evidences to support our conclusions that reduced gene dosage of *H4* contributes to defects in Cse4 sumoylation and reduced mislocalization of overexpressed Cse4 at peri-centromeric and non-centromeric regions, which in turn results in faster degradation of Cse4 and suppression of the *psh1Δ GALCSE4* SDL. The suppression of SDL by *hhf1*Δ and *hhf2*Δ is not limited to a *psh1*Δ *GALCSE4* background but is also observed in other mutants that exhibit *GALCSE4* associated SDL. Most importantly, our results with the *hhf1-20, cse4-102*, and *cse4-111* mutants, which are defective in the Cse4-H4 interaction, showed that the Cse4-H4 interaction is essential for non-centromeric association of Cse4. These studies are important from a clinical standpoint given the poor prognosis of CENP-A overexpressing cancers (Tomonaga *et al.* 2003; Amato *et al.* 2009; Li *et al.* 2011; McGovern *et al.* 2012; Sun *et al.* 2016; Zhang *et al.* 2016). Future studies with histone H4 and other mutants identified in our screen will provide insights into mechanisms that promote mislocalization of overexpressed Cse4 and how defects in these mechanisms may safeguard the cell from the lethal effect due to mislocalization of overexpressed Cse4 in mutants such as *psh1Δ*.

## ACKNOWLEDGEMENTS

We gratefully acknowledge Jennifer Gerton and Mitch Smith for reagents, Kathy McKinnon of the National Cancer Institute Vaccine Branch FACS Core for assistance with FACS analysis, Anthony Dawson for strain construction, and the members of the Basrai laboratory for helpful discussions and comments on the manuscript. MAB is supported by the NIH Intramural Research Program at the National Cancer Institute. This research was also supported by grants from the National Institutes of Health to CB and MC (R01HG005853) and from the Canadian Institute of Health Research to CB (FDN-143264). CB is a fellow in the Canadian Institute for Advanced Research (CIFAR, https://www.cifar.ca/) Fungal Kingdom: Threats and Opportunities. The funders had no role in study design, data collection and analysis, decision to publish, or preparation of the manuscript.

## CITATIONS

Allshire, R. C., and G. H. Karpen, 2008 Epigenetic regulation of centromeric chromatin: old dogs, new tricks? Nat Rev Genet 9: 923–937.

Amato, A., T. Schillaci, L. Lentini and A. Di Leonardo, 2009 CENPA overexpression promotes genome instability in pRb-depleted human cells. Mol Cancer 8: 119.

Athwal, R. K., M. P. Walkiewicz, S. Baek, S. Fu, M. Bui et al., 2015 CENP-A nucleosomes localize to transcription factor hotspots and subtelomeric sites in human cancer cells. Epigenetics Chromatin 8: 2.

Au, W. C., M. J. Crisp, S. Z. DeLuca, O. J. Rando and M. A. Basrai, 2008 Altered dosage and mislocalization of histone H3 and Cse4p lead to chromosome loss in Saccharomyces cerevisiae. Genetics 179: 263–275.

Au, W. C., A. R. Dawson, D. W. Rawson, S. B. Taylor, R. E. Baker et al., 2013 A Novel Role of the N-Terminus of Budding Yeast Histone H3 Variant Cse4 in Ubiquitin-Mediated Proteolysis. Genetics 194: 513–518.

Au, W. C., T. Zhang, P. K. Mishra, J. R. Eisenstatt, R. L. Walker et al., 2020 Skp, Cullin, F-box (SCF)-Met30 and SCF-Cdc4-Mediated Proteolysis of CENP-A Prevents Mislocalization of CENP-A for Chromosomal Stability in Budding Yeast. PLoS Genet 16: e1008597.

Biggins, S., 2013 The Composition, Functions, and Regulation of the Budding Yeast Kinetochore. Genetics 194: 817–846.

Boltengagen, M., A. Huang, A. Boltengagen, L. Trixl, H. Lindner et al., 2016 A novel role for the histone acetyltransferase Hat1 in the CENP-A/CID assembly pathway in Drosophila melanogaster. Nucleic Acids Res 44: 2145–2159.

Burrack, L. S., and J. Berman, 2012 Flexibility of centromere and kinetochore structures. Trends Genet 28: 204–212.

Camahort, R., B. Li, L. Florens, S. K. Swanson, M. P. Washburn et al., 2007 Scm3 is essential to recruit the histone H3 variant Cse4 to centromeres and to maintain a functional kinetochore. Mol Cell 26: 853–865.

Camahort, R., M. Shivaraju, M. Mattingly, B. Li, S. Nakanishi et al., 2009 Cse4 is part of an octameric nucleosome in budding yeast. Mol Cell 35: 794–805.

Chen, C. C., M. L. Dechassa, E. Bettini, M. B. Ledoux, C. Belisario et al., 2014 CAL1 is the Drosophila CENP-A assembly factor. J Cell Biol 204: 313–329.

Cheng, H., X. Bao, X. Gan, S. Luo and H. Rao, 2017 Multiple E3s promote the degradation of histone H3 variant Cse4. Sci Rep 7: 8565.

Cheng, H., X. Bao and H. Rao, 2016 The F-box Protein Rcy1 Is Involved in the Degradation of Histone H3 Variant Cse4 and Genome Maintenance. J Biol Chem 291: 10372–10377.

Chereji, R. V., J. Ocampo and D. J. Clark, 2017 MNase-Sensitive Complexes in Yeast: Nucleosomes and Non-histone Barriers. Mol Cell 65: 565–577 e563.

Choy, J. S., R. Acuna, W. C. Au and M. A. Basrai, 2011 A role for histone H4K16 hypoacetylation in Saccharomyces cerevisiae kinetochore function. Genetics 189: 11–21.

Choy, J. S., P. K. Mishra, W. C. Au and M. A. Basrai, 2012 Insights into assembly and regulation of centromeric chromatin in Saccharomyces cerevisiae. Biochim Biophys Acta 1819: 776–783.

Ciftci-Yilmaz, S., W. C. Au, P. K. Mishra, J. R. Eisenstatt, J. Chang et al., 2018 A Genome-Wide Screen Reveals a Role for the HIR Histone Chaperone Complex in Preventing Mislocalization of Budding Yeast CENP-A. Genetics 210: 203–218.

Cole, H. A., J. Ocampo, J. R. Iben, R. V. Chereji and D. J. Clark, 2014 Heavy transcription of yeast genes correlates with differential loss of histone H2B relative to H4 and queued RNA polymerases. Nucleic Acids Res 42: 12512–12522.

Collins, K. A., S. Furuyama and S. Biggins, 2004 Proteolysis contributes to the exclusive centromere localization of the yeast Cse4/CENP-A histone H3 variant. Curr Biol 14: 1968–1972.

Costanzo, M., B. VanderSluis, E. N. Koch, A. Baryshnikova, C. Pons et al., 2016 A global genetic interaction network maps a wiring diagram of cellular function. Science 353.

Cross, S. L., and M. M. Smith, 1988 Comparison of the structure and cell cycle expression of mRNAs encoded by two histone H3-H4 loci in Saccharomyces cerevisiae. Mol Cell Biol 8: 945–954.

Deyter, G. M., and S. Biggins, 2014 The FACT complex interacts with the E3 ubiquitin ligase Psh1 to prevent ectopic localization of CENP-A. Genes Dev 28: 1815–1826.

Deyter, G. M., E. M. Hildebrand, A. D. Barber and S. Biggins, 2017 Histone H4 Facilitates the Proteolysis of the Budding Yeast CENP-ACse4 Centromeric Histone Variant. Genetics 205: 113–124.

Eisenstatt, J. R., L. Boeckmann, W. C. Au, V. Garcia, L. Bursch et al., 2020 Dbf4-Dependent Kinase (DDK)-Mediated Proteolysis of CENP-A Prevents Mislocalization of CENP-A in Saccharomyces cerevisiae. G3 (Bethesda).

Fillingham, J., P. Kainth, J.-P. Lambert, H. van Bakel, K. Tsui et al., 2009 Two-color cell array screen reveals interdependent roles for histone chaperones and a chromatin boundary regulator in histone gene repression. Mol Cell 35: 340–351.

Foltz, D. R., L. E. Jansen, A. O. Bailey, J. R. Yates, 3rd, E. A. Bassett et al., 2009 Centromere-specific assembly of CENP-a nucleosomes is mediated by HJURP. Cell 137: 472–484.

Fujita, Y., T. Hayashi, T. Kiyomitsu, Y. Toyoda, A. Kokubu et al., 2007 Priming of centromere for CENP-A recruitment by human hMis18alpha, hMis18beta, and M18BP1. Dev Cell 12: 17–30.

Glowczewski, L., P. Yang, T. Kalashnikova, M. S. Santisteban and M. M. Smith, 2000 Histone-histone interactions and centromere function. Mol Cell Biol 20: 5700–5711.

Heun, P., S. Erhardt, M. D. Blower, S. Weiss, A. D. Skora et al., 2006 Mislocalization of the Drosophila centromere-specific histone CID promotes formation of functional ectopic kinetochores. Dev Cell 10: 303–315.

Hewawasam, G., M. Shivaraju, M. Mattingly, S. Venkatesh, S. Martin-Brown et al., 2010 Psh1 is an E3 ubiquitin ligase that targets the centromeric histone variant Cse4. Mol Cell 40: 444–454.

Hewawasam, G. S., K. Dhatchinamoorthy, M. Mattingly, C. Seidel and J. L. Gerton, 2018 Chromatin assembly factor-1 (CAF-1) chaperone regulates Cse4 deposition into chromatin in budding yeast. Nucleic Acids Res 46: 4440–4455.

Hewawasam, G. S., M. Mattingly, S. Venkatesh, Y. Zhang, L. Florens et al., 2014 Phosphorylation by casein kinase 2 facilitates Psh1 protein-assisted degradation of Cse4 protein. J Biol Chem 289: 29297–29309.

Hildebrand, E. M., and S. Biggins, 2016 Regulation of Budding Yeast CENP-A levels Prevents Misincorporation at Promoter Nucleosomes and Transcriptional Defects. PLoS Genet 12: e1005930.

Kastenmayer, J. P., L. Ni, A. Chu, L. E. Kitchen, W. C. Au et al., 2006 Functional genomics of genes with small open reading frames (sORFs) in S. cerevisiae. Genome Res 16: 365–373.

Kitagawa, K., and P. Hieter, 2001 Evolutionary concervation between budding yeast and human kinetochores. Nature Reviews Molecular Cellular Biology 2: 678–687.

Kurat, C. F., J. Recht, E. Radovani, T. Durbic, B. Andrews et al., 2014 Regulation of histone gene transcription in yeast. Cell Mol LIfe Sci 71: 599–613.

Lacoste, N., A. Woolfe, H. Tachiwana, A. V. Garea, T. Barth et al., 2014 Mislocalization of the centromeric histone variant CenH3/CENP-A in human cells depends on the chaperone DAXX. Mol Cell 53: 631–644.

Li, Y., Z. Zhu, S. Zhang, D. Yu, H. Yu et al., 2011 ShRNA-targeted centromere protein A inhibits hepatocellular carcinoma growth. PLoS One 6: e17794.

Maddox, P. S., K. D. Corbett and A. Desai, 2012 Structure, assembly and reading of centromeric chromatin. Curr Opin Genet Dev 22: 139–147.

McGovern, S. L., Y. Qi, L. Pusztai, W. F. Symmans and T. A. Buchholz, 2012 Centromere protein-A, an essential centromere protein, is a prognostic marker for relapse in estrogen receptor-positive breast cancer. Breast Cancer Res 14: R72.

McKinley, K. L., and I. M. Cheeseman, 2016 The molecular basis for centromere identity and function. Nat Rev Mol Cell Biol 17: 16–29.

Mishra, P. K., W. C. Au, J. S. Choy, P. H. Kuich, R. E. Baker et al., 2011 Misregulation of Scm3p/HJURP causes chromosome instability in Saccharomyces cerevisiae and human cells. PLoS Genet 7: e1002303.

Mizuguchi, G., H. Xiao, J. Wisniewski, M. M. Smith and C. Wu, 2007 Nonhistone Scm3 and histones CenH3-H4 assemble the core of centromere-specific nucleosomes. Cell 129: 1153–1164.

Moreno-Moreno, O., M. Torras-Llort and F. Azorin, 2006 Proteolysis restricts localization of CID, the centromere-specific histone H3 variant of Drosophila, to centromeres. Nucleic Acids Res 34: 6247–6255.

Ohkuni, K., R. Abdulle and K. Kitagawa, 2014 Degradation of centromeric histone H3 variant Cse4 requires the Fpr3 peptidyl-prolyl Cis-Trans isomerase. Genetics 196: 1041–1045.

Ohkuni, K., R. Levy-Myers, J. Warren, W. C. Au, Y. Takahashi et al., 2018 N-terminal Sumoylation of Centromeric Histone H3 Variant Cse4 Regulates Its Proteolysis To Prevent Mislocalization to Non-centromeric Chromatin. G3 (Bethesda) 8: 1215–1223.

Ohkuni, K., E. Suva, W. C. Au, R. L. Walker, R. Levy-Myers et al., 2020 Deposition of Centromeric Histone H3 Variant CENP-A/Cse4 into Chromatin Is Facilitated by Its C-Terminal Sumoylation. Genetics 214: 839–854.

Ohkuni, K., Y. Takahashi and M. A. Basrai, 2015 Protein purification technique that allows detection of sumoylation and ubiquitination of budding yeast kinetochore proteins Ndc10 and Ndc80. J Vis Exp: e52482.

Ohkuni, K., Y. Takahashi, A. Fulp, J. Lawrimore, W. C. Au et al., 2016 SUMO-Targeted Ubiquitin Ligase (STUbL) Slx5 regulates proteolysis of centromeric histone H3 variant Cse4 and prevents its mislocalization to euchromatin. Mol Biol Cell.

Pidoux, A. L., E. S. Choi, J. K. Abbott, X. Liu, A. Kagansky et al., 2009 Fission yeast Scm3: A CENP-A receptor required for integrity of subkinetochore chromatin. Mol Cell 33: 299–311.

Poch, O., and B. Winsor, 1997 Who’s Who among the Saccharomyces cerevisiae Actin-Related Proteins? A Classification and Nomenclature Proposal for a Large Family. Yeast 13: 1053–1058.

Prochasson, P., L. Florens, S. K. Swanson, M. P. Washburn and J. L. Workman, 2005 The HIR corepressor complex binds to nucleosomes generating a distinct protein/DNA complex resistant to remodeling by SWI/SNF. Genes Dev 19: 2534–2539.

Ranjitkar, P., M. O. Press, X. Yi, R. Baker, M. J. MacCoss et al., 2010 An E3 ubiquitin ligase prevents ectopic localization of the centromeric histone H3 variant via the centromere targeting domain. Mol Cell 40: 455–464.

Shen, X., G. Mizuguchi, A. Hamiche and C. Wu, 2000 A chromatin remodelling complex involved in transcription and DNA processing. Nature 406: 541–544.

Shen, X., R. Ranallo, E. Choi and C. Wu, 2003 Inolvement of Actin-Related Proteins in ATP-Dependent Chromatin Remodeling. Mol Cell 12: 147–155.

Shrestha, R. L., G. S. Ahn, M. I. Staples, K. M. Sathyan, T. S. Karpova et al., 2017 Mislocalization of centromeric histone H3 variant CENP-A contributes to chromosomal instability (CIN) in human cells. Oncotarget 8: 46781–46800.

Shuaib, M., K. Ouararhni, S. Dimitrov and A. Hamiche, 2010 HJURP binds CENP-A via a highly conserved N-terminal domain and mediates its deposition at centromeres. Proc Natl Acad Sci U S A 107: 1349–1354.

Smith, M. M., H. Yang, M. S. Santisteban, P. W. Boone, A. T. Goldstein et al., 1996 A Novel Histone H4 Mutant Defective in Nuclear Division and Mitotic Chromosome Transmission. Mol Cell Biol 16: 1017–1026.

Stoler, S., K. Rogers, S. Weitze, L. Morey, M. Fitzgerald-Hayes et al., 2007 Scm3, an essential Saccharomyces cerevisiae centromere protein required for G2/M progression and Cse4 localization. Proc Natl Acad Sci U S A 104: 10571–10576.

Sun, X., P. L. Clermont, W. Jiao, C. D. Helgason, P. W. Gout et al., 2016 Elevated expression of the centromere protein-A(CENP-A)-encoding gene as a prognostic and predictive biomarker in human cancers. Int J Cancer 139: 899–907.

Tomonaga, T., K. Matsushita, S. Yamaguchi, T. Oohashi, H. Shimada et al., 2003 Overexpression and mistargeting of centromere protein-A in human primary colorectal cancer. Cancer Res 63: 3511–3516.

Tosi, A., C. Haas, F. Herzog, A. Gilmozzi, O. Berninghausen et al., 2013 Structure and Subunit Topology of the INO80 Chromatin Remodeler and Its Nucleosome Complex. Cell 154: 1207–1219.

Verdaasdonk, J. S., and K. Bloom, 2011 Centromeres: unique chromatin structures that drive chromosome segregation. Nat Rev Mol Cell Biol 12: 320–332.

Williams, J. S., T. Hayashi, M. Yanagida and P. Russell, 2009 Fission yeast Scm3 mediates stable assembly of Cnp1/CENP-A into centromeric chromatin. Mol Cell 33: 287–298.

Zhang, W., J. H. Mao, W. Zhu, A. K. Jain, K. Liu et al., 2016 Centromere and kinetochore gene misexpression predicts cancer patient survival and response to radiotherapy and chemotherapy. Nat Commun 7: 12619.

Zhou, Z., H. Feng, B.-R. Zhuou, R. Ghirlando, K. Hu et al., 2011 Structural basis for recognition of centromere histone variant CenH3 by the chaperone Scm3. Nature 472: 234–237.

