## Supporting File S1 for "Reduced Gene Dosage of Histone H4 Prevents CENP-A Mislocalization in *Saccharomyces cerevisiae*"

Table S2: Yeast strains used in this study

| **Strain** | **Genotype** | **Parent Strain/Source** |
| --- | --- | --- |
| BY4741 | MAT**a** *his3∆1 leu2∆O met15∆O ura3∆O* | Open Biosystems |
| BY4742 | MAT**α** *his3∆1 leu2∆O met15∆O ura3∆O* | Open Biosystems |
| Y7092 | MAT**α** *his3∆1 leu2∆0 ura3∆0 met15∆0 lyp1∆ can1∆::STE2pr-SpHIS5* | Charlie Boone |
| YMB7290 | MAT**α** *cse4Δ::KanMX6 (pRB199=pRS316-6His-3HA-Cse4)* | (Ohkuni *et al.* 2016) |
| YMB8995 | MAT**α** *psh1*∆*::NAT^R^ his3∆1 leu2∆0 ura3∆0 met15∆0 lyp1∆ can1^::STE2pr-Sp_his5 lyp1^::STE3pr-LEU2* | Charlie Boone |
| YMB9034 | MAT**a** *psh1Δ::G418^R^ his3∆1 leu2∆O met15∆O ura3∆O* | BY4741 |
| YMB9802 | MAT**α** *his3∆1 leu2∆0 ura3∆0 met15∆0 lyp1∆ can1∆::STE2pr-SpHIS5 pMB433* | Y7092 |
| YMB9803 | MAT**α** *his3∆1 leu2∆0 ura3∆0 met15∆0 lyp1∆ can1∆::STE2pr-SpHIS5 pMB1458* | Y7092 |
| YMB10478 | MAT**α** *psh1*∆*::NAT^R^ his3∆1 leu2∆0 ura3∆0 met15∆0 lyp1∆ can1^::STE2pr-Sp_his5 lyp1^::STE3pr-LEU2 pMB433* | YMB8995 |
| YMB10479 | MAT**α** *psh1*∆*::NAT^R^ his3∆1 leu2∆0 ura3∆0 met15∆0 lyp1∆can1^::STE2pr-Sp_his5 lyp1^::STE3pr-LEU2 pMB1458* | YMB8995 |
| YMB10766 | MAT**a** *hhf1Δ::G418^R^ his3Δ1 leu2Δ0 met15Δ0 ura3Δ0* | Open Biosystems |
| YMB10767 | MAT**a** *hhf2Δ::G418^R^ his3Δ1 leu2Δ0 met15Δ0 ura3Δ0* | Open Biosystems |
| YMB10825 | MAT**a** *hhf1Δ::G418^R^ his3Δ1 leu2Δ0 met15Δ0 ura3Δ0 pMB433* | YMB10766 x YMB9802 |
| YMB11166 | MAT**a** *hhf2Δ::G418^R^ his3Δ1 leu2Δ0 met15Δ0 ura3Δ0* *pMB433* | YMB10767 x YMB9802 |
| YMB10937 | MAT**a** *hhf1∆::G418^R^ his3Δ1 leu2Δ0 met15Δ0 ura3Δ0 pMB1458* | YMB10766 x YMB9803 |
| YMB10938 | MAT**a** *hhf2∆::G418^R^ his3Δ1 leu2Δ0 met15Δ0 ura3Δ0 pMB1458* | YMB10767 x YMB9803 |
| YMB10821 | MAT**a** *psh1Δ::NAT^R^ hhf1Δ::G418^R^ pMB433* | YMB10766 x YMB10478 |
| YMB10822 | MAT**a** *psh1Δ::NAT^R^ hhf1Δ::G418^R^ pMB1458* | YMB10766 x YMB10479 |
| YMB10823 | MAT**a** *psh1Δ::NAT^R^ hhf2Δ::G418^R^ GAL-pMB433* | YMB10767 x YMB10478 |
| YMB10824 | MAT**a** *psh1Δ::NAT^R^ hhf2Δ::G418^R^ pMB1458* | YMB10767 x YMB10479 |
| YMB8994 | MAT**α** *slx5*∆*::NAT^R^ his3∆1 leu2∆0 ura3∆0 met15∆0 lyp1∆ can1^::STE2pr-Sp_his5 lyp1^::STE3pr-LEU2* | Charlie Boone |
| YMB10963 | MAT**α** *slx5*∆*::NAT^R^ his3∆1 leu2∆0 ura3∆0 met15∆0 lyp1∆ can1^::STE2pr-Sp_his5 lyp1^::STE3pr-LEU2 pMB1458* | YMB8994 |
| YMB11046 | MAT**a** *slx5*∆*::NAT^R^ hhf1::G418^R^ pMB1458* | YMB10963 x YMB10766 |
| YMB11047 | MAT**a** *slx5*∆*::NAT^R^ hhf2::G418^R^ pMB1458* | YMB10963 x YMB10767 |
| SN2176 | MAT**α** doa*1*∆*::NAT^R^ his3∆1 leu2∆0 ura3∆0 met15∆0 lyp1∆ can1^::STE2pr-Sp_his5 lyp1^::STE3pr-LEU2* | Charlie Boone |
| YMB11032 | MAT**α** doa*1*∆*::NAT^R^ his3∆1 leu2∆0 ura3∆0 met15∆0 lyp1∆ can1^::STE2pr-Sp_his5 lyp1^::STE3pr-LEU2 pMB1458* | SN2176 |
| YMB11050 | MAT**a** *doa1Δ::NAT^R^ hhf1Δ::G418^R^ pMB1458* | YMB11032 x YMB10766 |
| YMB11053 | MAT**a** *doa1Δ::NAT^R^ hhf2Δ::G418^R^ pMB1458* | YMB11032 x YMB10767 |
| YMB8785 | MAT**α** *hir2*∆*::NAT^R^ his3∆1 leu2∆O lys2∆0 ura3∆O* | BY4742 |
| YMB8332 | MAT**a** *hir2∆::NAT^R^ his3∆1 leu2∆O met15∆O ura3∆O pMB1458* | BY4741 |
| YMB11105 | MAT**α** *hir2*∆*::NAT^R^ hhf1*∆*::G418^R^ pMB1458* | YMB8785 x YMB10937 |
| YMB11107 | MAT**a** *hir2∆::NAT^R^ hhf2∆::G418^R^ pMB1458* | YMB8785 x YMB10938 |
| YMB8933 | MAT**α** *cdc4-1::NAT^R^ his3∆1 leu2∆0 ura3∆0 met15∆0 lyp1∆ can1^::STE2pr-Sp_his5 lyp1^::STE3pr-LEU2* | Charlie Boone |
| YMB9756 | MAT**α** *cdc4-1::NAT^R^ his3∆1 leu2∆0 ura3∆0 met15∆0 lyp1∆ can1^::STE2pr-Sp_his5 lyp1^::STE3pr-LEU2 pMB1458* | YMB8933 |
| YMB11051 | MAT**a** *cdc4-1::NAT^R^ hhf1Δ::G418^R^ pMB1458* | YMB9756 x YMB10766 |
| YMB11054 | MAT**a** *cdc4-1::NAT^R^ hhf2Δ::G418^R^ pMB1458* | YMB9756 x YMB10767 |
| tsa131 | MAT**α** *cdc7-4::NAT^R^ his3∆1 leu2∆0 ura3∆0 met15∆0 lyp1∆ can1^::STE2pr-Sp_his5 lyp1^::STE3pr-LEU2* | Charlie Boone |
| YMB9760 | MAT**α** *cdc7-4::NAT^R^ his3∆1 leu2∆0 ura3∆0 met15∆0 lyp1∆ can1^::STE2pr-Sp_his5 lyp1^::STE3pr-LEU2 pMB1458* | tsa131 |
| YMB11052 | MAT**a** *cdc7-4::NAT^R^ hhf1Δ::G418^R^ pMB1458* | YMB9760 x YMB10766 |
| YMB11055 | MAT**a** *cdc7-4::NAT^R^ hhf2Δ::G418^R^ pMB1458* | YMB9760 x YMB10767 |
| YMB11178 | MAT**a** *hta1Δ::G418^R^ his3Δ1 leu2Δ0 met15Δ0 ura3Δ0* | Open Biosystems |
| YMB11258 | MAT**a** *hta1Δ::G418^R^ pMB433* | YMB11178 x YMB10478 |
| YMB11262 | MAT**a** *hta1Δ::G418^R^ pMB1458* | YMB11178 x YMB10479 |
| YMB11260 | MAT**a** *psh1Δ::NAT^R^ hta1Δ::G418^R^ pMB433* | YMB11178 x YMB10478 |
| YMB11264 | MAT**a** *psh1Δ::NAT^R^ hta1Δ::G418^R^ pMB1458* | YMB11178 x YMB10479 |
| YMB11179 | MAT**a** *hta2Δ::G418^R^ his3Δ1 leu2Δ0 met15Δ0 ura3Δ0* | Open Biosystems |
| YMB11266 | MAT**a** *hta2Δ::G418^R^ pMB433* | YMB11179 x YMB10478 |
| YMB11270 | MAT**a** *hta2Δ::G418^R^ pMB1458* | YMB11179 x YMB10479 |
| YMB11268 | MAT**a** *psh1Δ::NAT^R^ hta2Δ::G418^R^ pMB433* | YMB11179 x YMB10478 |
| YMB11272 | MAT**a** *psh1Δ::NAT^R^ hta2Δ::G418^R^ pMB1458* | YMB11179 x YMB10479 |
| YMB11180 | MAT**a** *hht1Δ::G418^R^ his3Δ1 leu2Δ0 met15Δ0 ura3Δ0* | Open Biosystems |
| YMB11274 | MAT**a** *hht1Δ::G418^R^ pMB433* | YMB11180 x YMB10478 |
| YMB11278 | MAT**a** *hht1Δ::G418^R^ pMB1458* | YMB11180 x YMB10479 |
| YMB11276 | MAT**a** *psh1**Δ::NAT^R^ hht1Δ::G418^R^ pMB433* | YMB11180 x YMB10478 |
| YMB11280 | MAT**a** *psh1Δ::NAT^R^ hht1Δ::G418^R^ pMB1458* | YMB11180 x YMB10479 |
| YMB11181 | MAT**a** *hht2Δ::G418^R^ his3Δ1 leu2Δ0 met15Δ0 ura3Δ0* | Open Biosystems |
| YMB11282 | MAT**a** *hht2Δ::G418^R^ pMB433* | YMB11181 x YMB10478 |
| YMB11286 | MAT**a** *hht2Δ::G418^R^ pMB1458* | YMB11181 x YMB10479 |
| YMB11284 | MAT**a** *psh1Δ::NAT^R^ hht2Δ::G418^R^ pMB433* | YMB11181 x YMB10478 |
| YMB11288 | MAT**a** *psh1Δ::NAT^R^ hht2Δ::G418^R^ pMB1458* | YMB11181 x YMB10479 |
| JG1689 | MAT**a** *pGAL1-10-3HA-SCM3::TRP1 <pSB17 (empty vector) URA3>* | (Hewawasam *et al.* 2018) |
| JG1690 | MAT**a** *pGAL1-10-3HA-SCM3::TRP1 <pSB873 (Cu-CSE4) URA3>* | (Hewawasam *et al.* 2018) |
| YMB11252 | MAT**a** *hhf2Δ::G418^R^ pGAL1-10-3HA-SCM3::TRP1 <pSB17 (empty vector) URA3>* | JG1689 |
| YMB11253 | MAT**a** *hhf2Δ::G418^R^ pGAL1-10-3HA-SCM3::TRP1 <pSB873 (Cu-CSE4) URA3>* | JG1690 |
| MSY559 | *MAT***a** *leu2,3-112 ura3-52 lys2*Δ*200 HHT1HHF1* Δ(*HHT2-HHF2*) | (Glowczewski *et al.* 2000) |
| MSY535 | *MAT***a** *leu2,3-112 ura3-52 lys2*Δ*200 HHT1 hhf1-10* Δ(*HHT1-HHF1*) Δ(*HHT2-HHF2*) | (Glowczewski *et al.* 2000) |
| MSY534 | *MAT***a** *leu2,3-112 ura3-52 lys2*Δ*200 HHT1 hhf1-20* Δ(*HHT1-HHF1*) Δ(*HHT2-HHF2*) | (Glowczewski *et al.* 2000) |
| YMB11346 | *MAT***a** *leu2,3-112 ura3-52 lys2*Δ*200 HHT1HHF1* Δ(*HHT2-HHF2*) | MSY559 |
| YMB11347 | *MAT***a** *leu2,3-112 ura3-52 lys2*Δ*200 HHT1 hhf1-10* Δ(*HHT1-HHF1*) Δ(*HHT2-HHF2*) *psh1Δ* | MSY535 |
| YMB11348 | *MAT***a** *leu2,3-112 ura3-52 lys2*Δ*200 HHT1 hhf1-20* Δ(*HHT1-HHF1*) Δ(*HHT2-HHF2*) *psh1Δ* | MSY534 |

Table S3: Plasmids used in this study.

| **Strain** | **Promoter, Gene, and Marker** | **Source** |
| --- | --- | --- |
| pRS415 | *CEN LEU2* | (Sikorski and Hieter 1989) |
| pMB1725 | *CEN 6His-3HA-CSE4 LEU2* | This Study |
| pMB1935 | *CEN 6His-3HA-cse^D217A^ LEU2* | This Study |
| pMB1936 | *CEN 6His-3HA-cse^D217E^ LEU2* | This Study |
| pRS425 | *2µ LEU2* | (Christianson *et al.* 1992) |
| pMB433 | *2µ pGAL1 URA3* | (Mumberg *et al.* 1994) |
| pMB1458 | *2µ pGAL1-6His-3HA-CSE4 URA3* | (Au *et al.* 2013) |
| pMB1892 | *2µ pGAL1-3HA-cse4^16KR^ URA3* | (Au *et al.* 2020) |
| pMB1928 | *MoBy 2µ HHF1 LEU2* | This Study |
| pMB1929 | *MoBy 2µ HHF2 LEU2* | This Study |
| pYES2 | *2µ pGAL URA3* | (Boeckmann *et al.* 2013) |
| pMB1344 | *2µ* pYES2-*pGAL-8His-HA-cse4^16KR^* *URA3* | (Ohkuni *et al.* 2016) |
| pMB1345 | *2µ* pYES2-*pGAL-8His-HA-CSE4* *URA3* | (Ohkuni *et al.* 2016) |
| pMB1766 | *2µ* pYES2-*pGAL-8His-HA-cse4 Y193A* *URA3* | This Study |
| pMB1787 | *2µ* pYES2-*pGAL-8His-HA-cse4 Y193F* *URA3* | This Study |
| pMB1910 | *2µ* pYES2-*pGAL-8His-HA-cse4 D217A* *URA3* | This Study |
| pMB1920 | *2µ* pYES2-*pGAL-8His-HA-cse4 D217E* *URA3* | This Study |
| pMB1984 | *2µ* pYES2-*pGAL-8His-HA-cse4-102* *URA3* | This Study |
| pMB1985 | *2µ* pYES2-*pGAL-8His-HA-cse4-107^MB^ URA3* | This Study |
| pMB1986 | *2µ* pYES2-*pGAL-8His-HA-cse4-108* *URA3* | This Study |
| pMB1987 | *2µ* pYES2-*pGAL-8His-HA-cse4-110* *URA3* | This Study |
| pMB1988 | *2µ* pYES2-*pGAL-8His-HA-cse4-111* *URA3* | This Study |

Table S4: Primers used in this study.

| **Region** | **Primer** | **Sequence** | **Source** |
| --- | --- | --- | --- |
| *ACT1* For | OMB444 | ACAACGAATTGAGAGTTGCCCCAG | (Dimova *et al.* 1999) |
| *ACT1* Rev | OMB445 | AATGGCGTGAGGTAGAGAGAAACC | (Dimova *et al.* 1999) |
| *SAP4* For | SB3735 | ACAGCACAACACGCTTACCA | (Hildebrand and Biggins 2016) |
| *SAP4* Rev | SB3736 | CCAGCCCTAAATCCCCTAAA | (Hildebrand and Biggins 2016) |
| *RDS1* For | SB4768 | GACCCGTGCAGATCACTATTACA | (Hildebrand and Biggins 2016) |
| *RDS1* Rev | SB4769 | GCAGTTTATCACATTTCCGTTTG | (Hildebrand and Biggins 2016) |
| *SLP1* For | SB3781 | TCCTAGGTTATCTCATCGGTACT | (Hildebrand and Biggins 2016) |
| *SLP1* Rev | SB3782 | ACTATATCCATTGCGTCCTTTCT | (Hildebrand and Biggins 2016) |
| *PHO5* For | OMB3282 | CCCATTTGGGATAAGGGTAAAC | (Hewawasam *et al.* 2018) |
| *PHO5* Rev | OMB3283 | GATGAAGCCATACTAACCTCGA | (Hewawasam *et al.* 2018) |
| *FIG4*/*LEM3* For | OMB3274 | ACCAGAACGGCAGACAAAGT | (Ohkuni *et al.* 2020) |
| *FIG3*/*LEM3* Rev | OMB3275 | TGAACGTGCTGCAATAAACC | (Ohkuni *et al.* 2020) |
| *UGA3*/*UGX2* For | OMB3298 | CCCCCTGGGTGTCTTAAATTAT | (Ohkuni *et al.* 2020) |
| *UGA3*/*UGX2* Rev | OMB3298 | TTCTATGTCTCAACGTTAGCATTTC | (Ohkuni *et al.* 2020) |
| *COQ3* For | OMB3276 | CCTTTCATAATGTATTATCACCCTTT | (Ohkuni *et al.* 2020) |
| *COQ3* Rev | OMB3277 | AGACTTCGCTGTACCTGTTTCC | (Ohkuni *et al.* 2020) |
| *GUP2* For | OMB3278 | CGATGGTATTGATGCACCTG | (Ohkuni *et al.* 2020) |
| *GUP2* Rev | OMB3279 | GACTGTTAACCACCCCCAAA | (Ohkuni *et al.* 2020) |
| peri*CEN3* L3 For | OMB1684 | GCCATACCATGCTTTGTTATCGTC | (Choy *et al.* 2011) |
| peri*CEN3* L3 Rev | OMB1685 | TATTATGCTCCCCTGGATTTTATGCG | (Choy *et al.* 2011) |
| peri*CEN3* L2 For | OMB1686 | TCATCTTTGAAAAGTTCATCAAGG | (Choy *et al.* 2011) |
| peri*CEN3* L2 Rev | OMB1687 | GATAACAAAGCATGGTATGGCG | (Choy *et al.* 2011) |
| peri*CEN3* L1 For | OMB1688 | ATATTGTTTGGCGCTGATCGCC | (Choy *et al.* 2011) |
| peri*CEN3* L1 Rev | OMB1689 | TTGATGAACTTTTCAAAGATGAC | (Choy *et al.* 2011) |
| *CEN3* For | OMB244 | GATCAGCGCCAAACAATATGG | (Choy *et al.* 2011) |
| *CEN3* Rev | OMB245 | AACTTCCACCAGTAAACGTTTC | (Choy *et al.* 2011) |
| peri*CEN3* R1 For | OMB1692 | TTTACTGGTGGAAGTTTTGCTCA | (Choy *et al.* 2011) |
| peri*CEN3* R1 Rev | OMB1693 | GTCAACGAGTCCTCTCTGGCTA | (Choy *et al.* 2011) |
| peri*CEN3* R2 For | OMB1694 | GAGAGGACTCGTTGACGTAGAA | (Choy *et al.* 2011) |
| peri*CEN3* R2 Rev | OMB1695 | GAATATGATAATGGTTACACCAGTAGG | (Choy *et al.* 2011) |
| peri*CEN3* R3 For | OMB1696 | TGTAACCATTATCATATTCATGAC | (Choy *et al.* 2011) |
| peri*CEN3* R3 Rev | OMB1697 | GATTTAATGCACGTTATGTTTCG | (Choy *et al.* 2011) |

CITATIONS

Au, W. C., A. R. Dawson, D. W. Rawson, S. B. Taylor, R. E. Baker *et al.*, 2013 A Novel Role of the N-Terminus of Budding Yeast Histone H3 Variant Cse4 in Ubiquitin-Mediated Proteolysis. Genetics 194**:** 513-518.

Au, W. C., T. Zhang, P. K. Mishra, J. R. Eisenstatt, R. L. Walker *et al.*, 2020 Skp, Cullin, F-box (SCF)-Met30 and SCF-Cdc4-Mediated Proteolysis of CENP-A Prevents Mislocalization of CENP-A for Chromosomal Stability in Budding Yeast. PLoS Genet 16**:** e1008597.

Boeckmann, L., Y. Takahashi, W. C. Au, P. K. Mishra, J. S. Choy *et al.*, 2013 Phosphorylation of centromeric histone H3 variant regulates chromosome segregation in Saccharomyces cerevisiae. Mol Biol Cell 24**:** 2034-2044.

Choy, J. S., R. Acuna, W. C. Au and M. A. Basrai, 2011 A role for histone H4K16 hypoacetylation in Saccharomyces cerevisiae kinetochore function. Genetics 189**:** 11-21.

Christianson, T. W., R. S. Sikorski, M. Dante, J. H. Shero and P. Hieter, 1992 Multifunctional yeast high-copy-number shuttle vectors. Gene 110**:** 119-122.

Dimova, D., Z. Nackerdien, S. Furgeson, S. Eguchi and M. A. Osley, 1999 A role for transcriptional repressors in targeting the yeast Swi/Snf complex. Mol Cell 4**:** 75-83.

Glowczewski, L., P. Yang, T. Kalashnikova, M. S. Santisteban and M. M. Smith, 2000 Histone-histone interactions and centromere function. Mol Cell Biol 20**:** 5700-5711.

Hewawasam, G. S., K. Dhatchinamoorthy, M. Mattingly, C. Seidel and J. L. Gerton, 2018 Chromatin assembly factor-1 (CAF-1) chaperone regulates Cse4 deposition into chromatin in budding yeast. Nucleic Acids Res 46**:** 4440-4455.

Hildebrand, E. M., and S. Biggins, 2016 Regulation of Budding Yeast CENP-A levels Prevents Misincorporation at Promoter Nucleosomes and Transcriptional Defects. PLoS Genet 12**:** e1005930.

Mumberg, D., R. Muller and M. Funk, 1994 Regulatable promoters of Saccharomyces cerevisiae: comparison of transcriptional activity and their use for heterologous expression. Nucleic Acids Res 22**:** 5767-5768.

Ohkuni, K., E. Suva, W. C. Au, R. L. Walker, R. Levy-Myers *et al.*, 2020 Deposition of Centromeric Histone H3 Variant CENP-A/Cse4 into Chromatin Is Facilitated by Its C-Terminal Sumoylation. Genetics 214**:** 839-854.

Ohkuni, K., Y. Takahashi, A. Fulp, J. Lawrimore, W. C. Au *et al.*, 2016 SUMO-Targeted Ubiquitin Ligase (STUbL) Slx5 regulates proteolysis of centromeric histone H3 variant Cse4 and prevents its mislocalization to euchromatin. Mol Biol Cell.

Sikorski, R. S., and P. Hieter, 1989 A System of Shuttle Vectors and Yeast Host Strains Designed for Efficient Manipulation of DNA in *Saccharomyces cerevisiae*. Genetics 122**:** 19-27.
