## Supplementary figures and images for "Reduced Gene Dosage of Histone H4 Prevents CENP-A Mislocalization in *Saccharomyces cerevisiae*"

### Supporting Figure S1

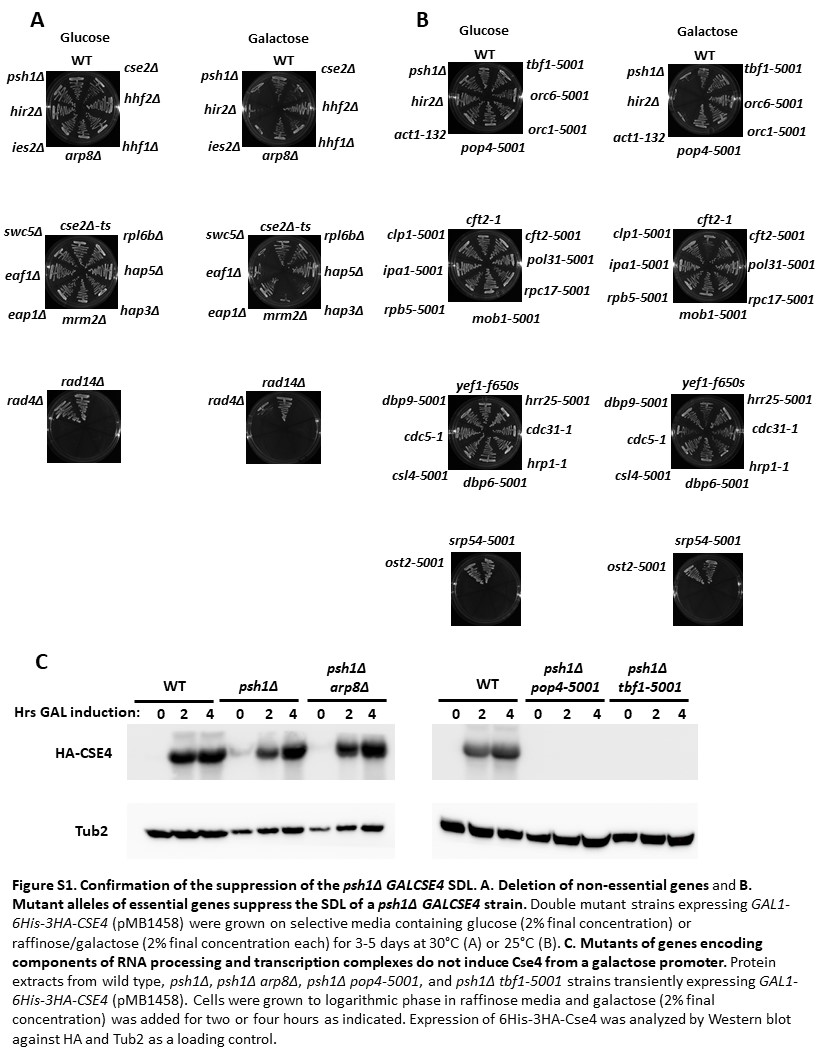

### Supporting figure S2

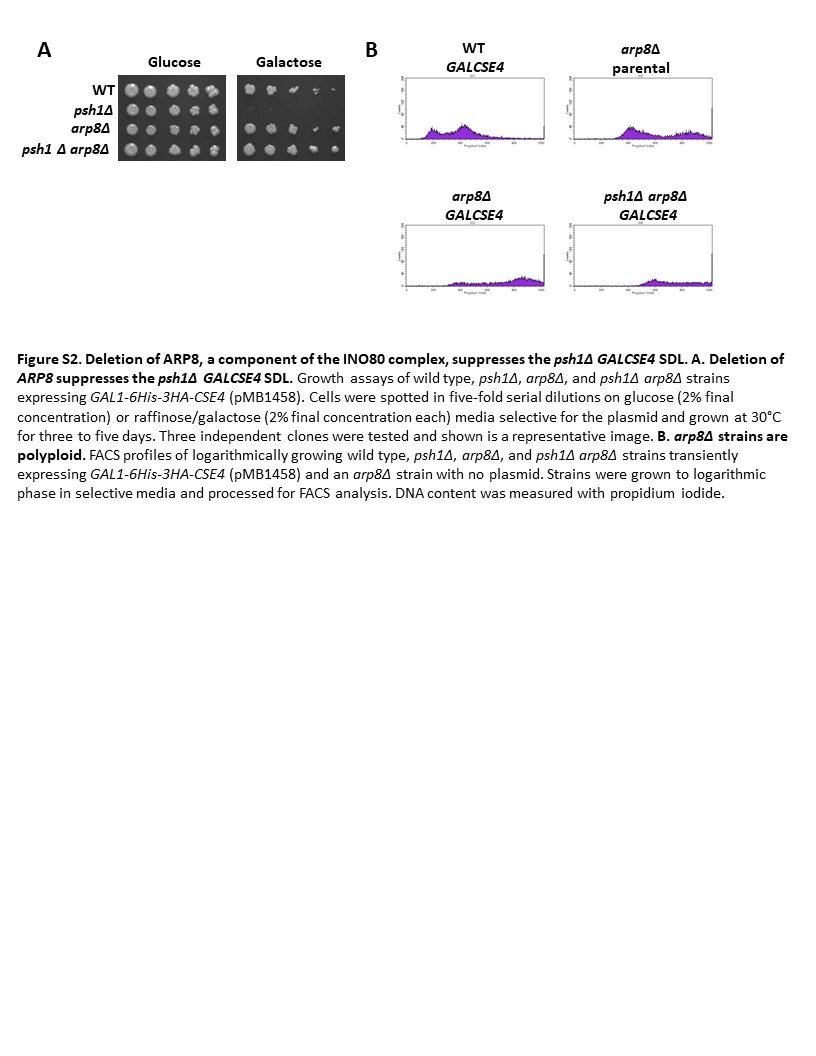

### Supporting Figure S3

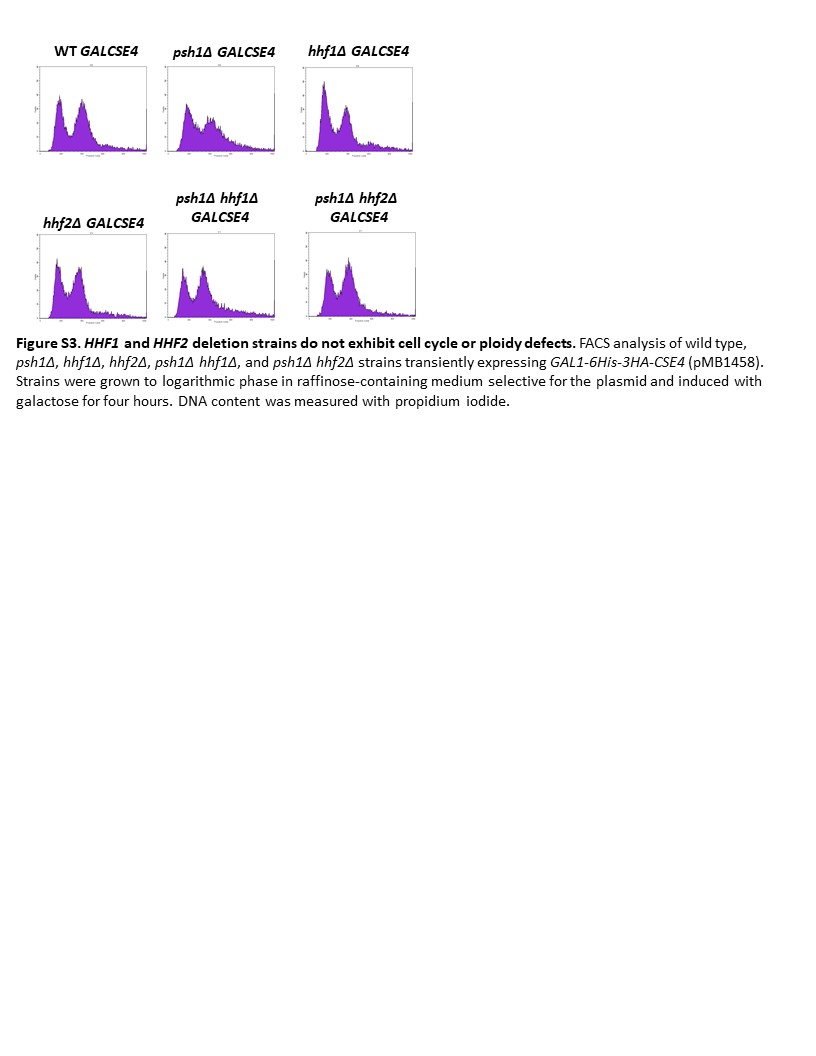

### Supporting Figure S4

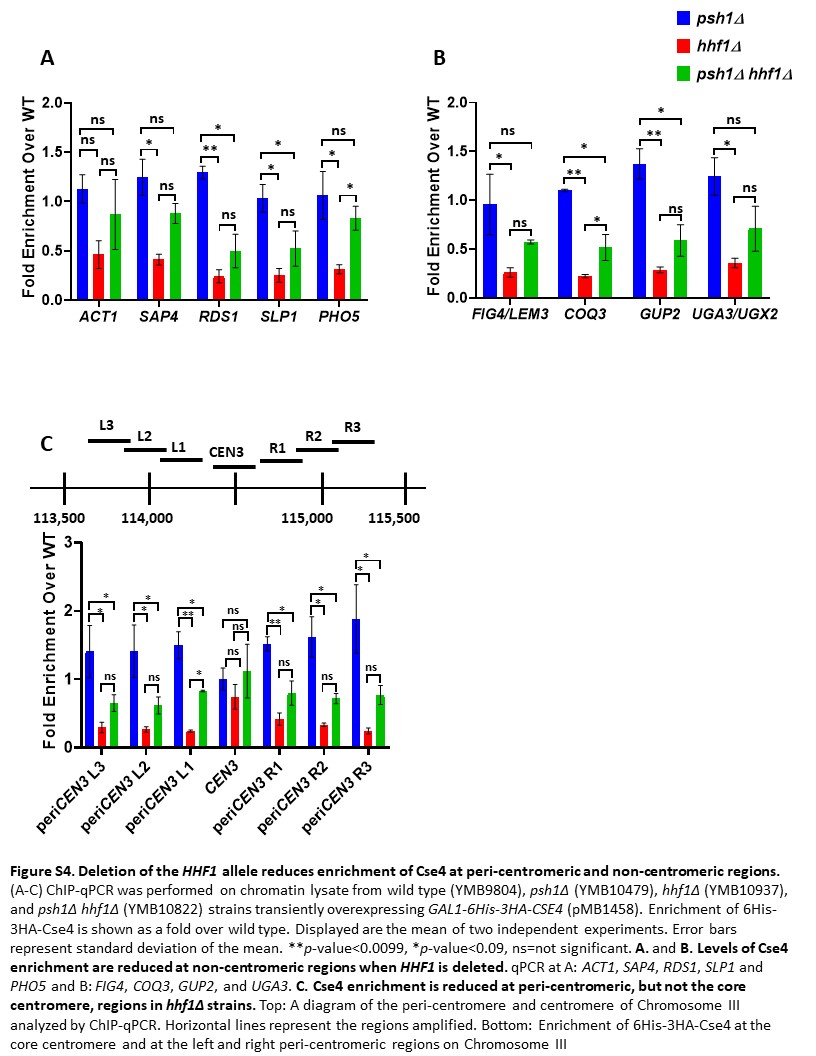

### Supporting Figure S5

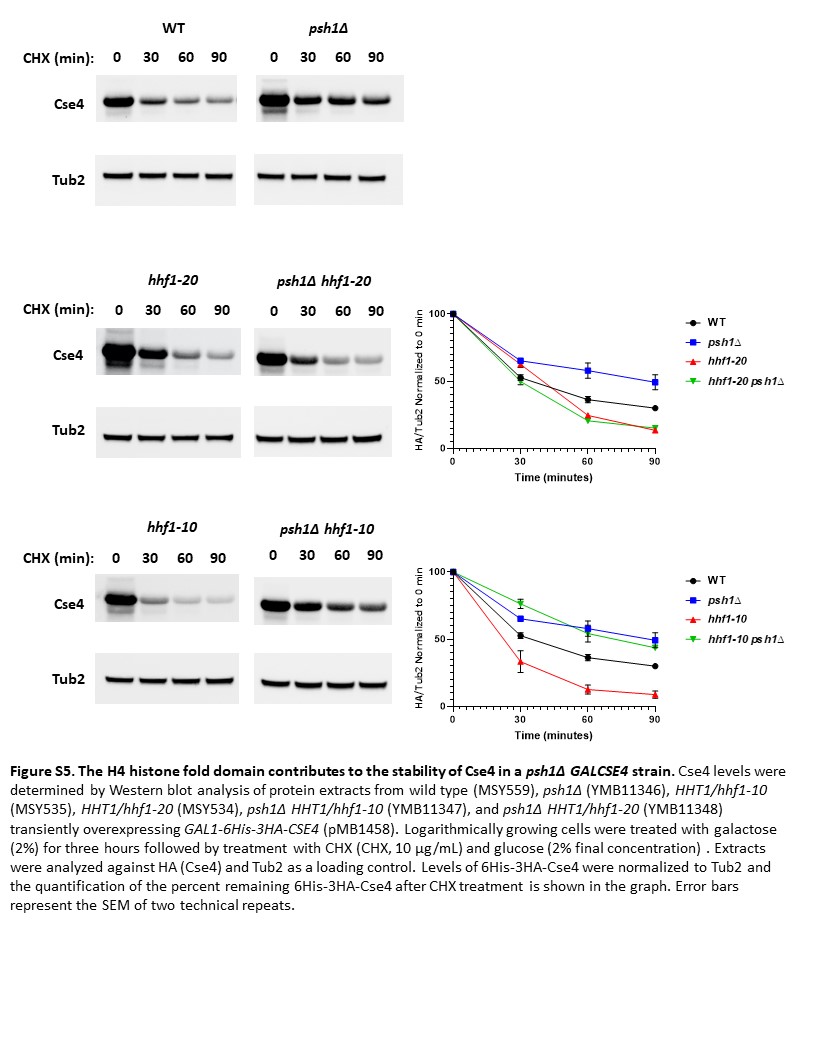

### Supporting Figure S6

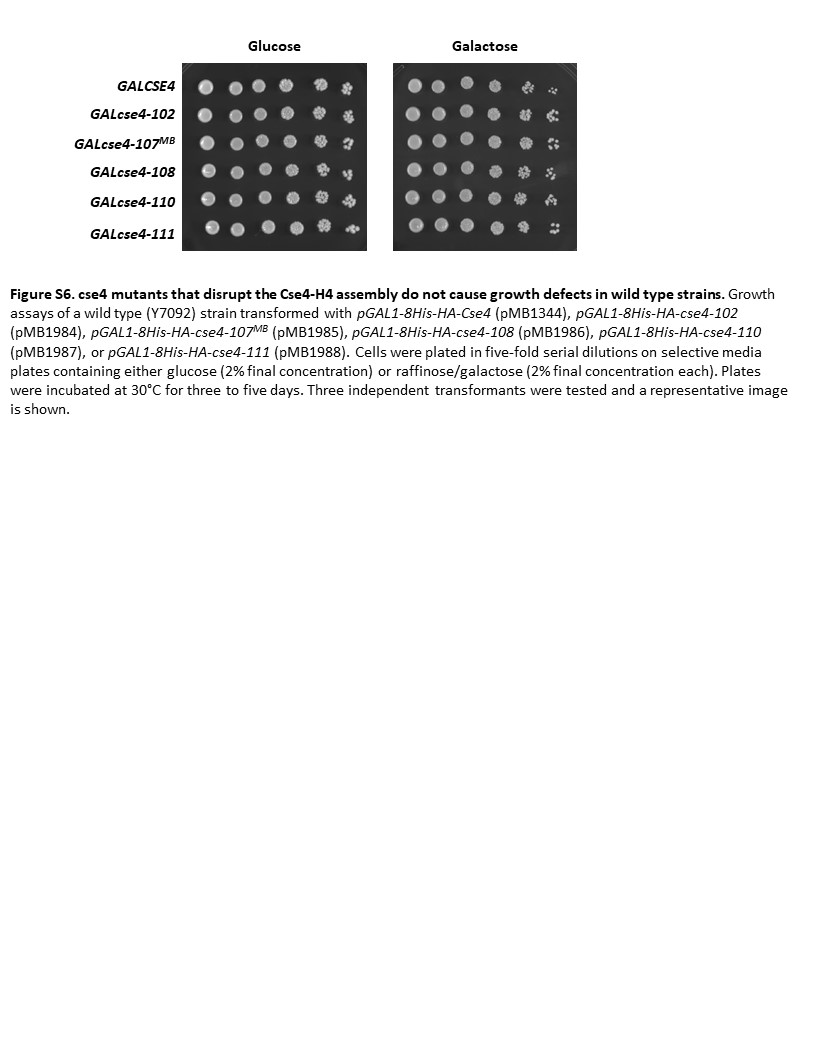

### Supporting Figure S7

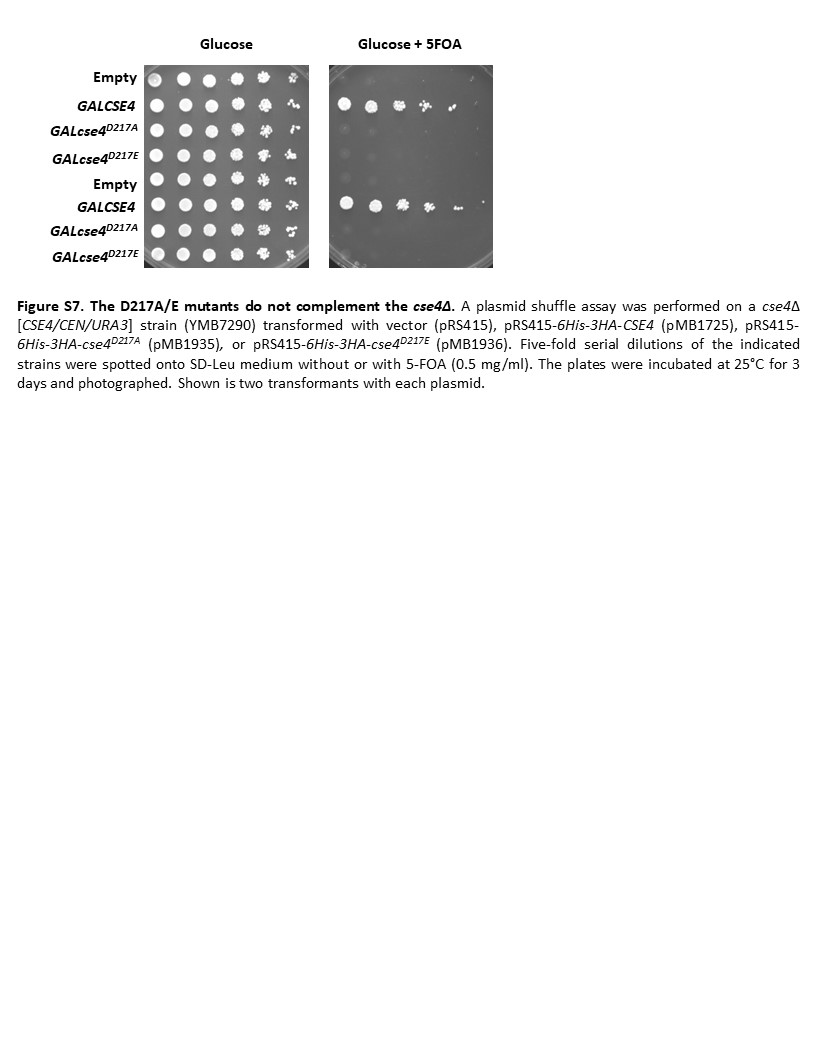
